# Therapeutic benefits of maintaining CDK4/6 inhibitors and incorporating CDK2 inhibitors beyond progression in breast cancer

**DOI:** 10.1101/2024.11.11.623139

**Authors:** Jessica Armand, Sungsoo Kim, Kibum Kim, Eugene Son, Minah Kim, Kevin Kalinsky, Hee Won Yang

**Affiliations:** Department of Pathology and Cell Biology, Columbia University Irving Medical Center, New York, NY, 10032, USA; Herbert Irving Comprehensive Cancer Center, Columbia University Irving Medical Center, New York, NY, 10032, USA; Winship Cancer Institute at Emory University, Department of Hematology and Medical Oncology, Atlanta, GA

## Abstract

CDK4/6 inhibitors (CDK4/6i) with endocrine therapy are standard for hormone receptor-positive (HR^+^) metastatic breast cancer. However, most patients eventually develop resistance and discontinue treatment, and there is currently no consensus on effective second-line strategies. Here, we demonstrate that maintaining CDK4/6i therapy, either alone or in combination with CDK2 inhibitors (CDK2i), slows the growth of drug-resistant HR^+^ models by prolonging G1 progression. Mechanistically, sustained CDK4/6 blockade in drug-resistant cells reduces E2F transcription and delays G1/S via a non-canonical, post-translational regulation of retinoblastoma protein (Rb). Durable suppression of both CDK2 activity and growth of drug-resistant cells requires co-administration of CDK2i with CDK4/6i. Moreover, cyclin E overexpression drives resistance to the combination of CDK4/6i and CDK2i. These findings elucidate how continued CDK4/6 blockade constrains resistant tumors and support clinical strategies that maintain CDK4/6i while selectively incorporating CDK2i to overcome resistance.

## Introduction

Metastatic breast cancer remains a leading cause of cancer-related mortality in women globally (1,2). A key dysregulation in breast cancer involves the overactivation of cyclin-dependent kinases 4 and 6 (CDK4/6) (3–6). Active CDK4/6 phosphorylates the retinoblastoma protein (Rb), a crucial regulator that prevents cell-cycle initiation by sequestering E2F transcription factors (7). Rb phosphorylation results in the release of E2F, thereby promoting CDK2 activation and cell proliferation (8,9). Understanding this mechanism has driven significant advancements in therapeutic strategies, particularly for hormone receptor-positive (HR^+^)/human epidermal growth factor receptor 2-negative (HER2^-^) breast cancer, which constitutes approximately 70% of breast cancer cases (10). The current standard first-line treatment for HR^+^/HER2^-^ metastatic breast cancer is a combination of CDK4/6 inhibitors (CDK4/6i) and endocrine therapy (ET) (4,5,11). Although this strategy has significantly improved patient outcomes, resistance remains a major challenge: approximately 30% of patients develop resistance within two years, and the majority ultimately relapse (12,13). Upon disease progression, CDK4/6i therapy is typically discontinued, often resulting in aggressive tumor regrowth and a lack of effective second-line treatment options.

While continuing ET after disease progression confers clinical benefit (14–18), the value of maintaining CDK4/6i treatment beyond progression remains elusive. Multiple ongoing and completed trials have evaluated continuing the FDA-approved CDK4/6 inhibitors, palbociclib, abemaciclib, and ribociclib, after progression (19–23). MAINTAIN and postMONARCH trials reported significantly improved progression-free survival (PFS) with continued CDK4/6i plus ET (21,23). Retrospective analyses likewise suggest that second-line CDK4/6i plus ET can outperform chemotherapy or ET alone in several settings (24,25). Safety appears comparable or better than first-line trials, with no new toxicities and grade 3–4 events largely confined to expected hematologic adverse effects. In contrast, the PACE and PALMIRA trials did not demonstrate a PFS benefit for continuing CDK4/6i plus ET after progression (19,20). Overall, the evidence for CDK4/6i continuation beyond progression is encouraging but mixed, underscoring the need for mechanistic studies of CDK4/6i-resistant proliferation to guide post-progression treatment strategies for HR^+^/HER2^-^ breast cancer.

Like CDK4/6, CDK2 also phosphorylates Rb before the G1/S transition (8,26). The relevance of CDK2 activation in CDK4/6i-resistant tumors highlights the potential of targeting CDK2 to overcome CDK4/6i resistance (27–34). Several clinical trials are currently evaluating CDK2 inhibitors (CDK2i) in patients who have progressed on CDK4/6i-based therapy, either as monotherapy or in combination with CDK4/6i, with or without ET (NCT05252416, NCT05735080). Determining the optimal strategy for incorporating CDK2i in treating HR^+^/HER2^-^breast cancer that has developed drug resistance remains a critical question.

This study shows that, even in drug-resistant cells that remain proliferative, continued CDK4/6 inhibition results in ineffective Rb inactivation, thereby slowing E2F activation kinetics and prolonging the G1 phase. These data suggest that overall survival, rather than PFS, may serve as a more appropriate clinical trial endpoint. Moreover, our work highlights the therapeutic potential of combining CDK2i with CDK4/6i as an effective second-line strategy. Finally, we identify cyclin E overexpression as a key driver of resistance to this combination, offering mechanistic guidance for overcoming therapy resistance.

## Results

### Maintaining CDK4/6i treatment attenuates the growth of drug-resistant cells by extending G1-phase progression

To evaluate the impact of continued CDK4/6i treatment in drug-resistant settings, we employed HR^+^/HER2^-^ breast cancer cell lines (MCF-7, T47D, and CAMA-1) and a triple-negative breast cancer cell line (MDA-MB-231), all of which have an intact Rb/E2F pathway. These cells were chronically exposed to palbociclib for over a month to induce drug resistance (Figure 1A). We confirmed increased half-maximal inhibitory concentrations (IC50) for palbociclib in drug-resistant cells compared to parental cells (Figure S1A). We maintained or withdrew CDK4/6i treatment in drug-resistant cells and monitored their cumulative proliferation alongside that of parental cells. Drug-resistant cells maintained on CDK4/6i exhibited significantly slower growth than those under other conditions (Figure 1B). In contrast, resistant cells withdrawn from treatment grew at rates comparable to parental cells. Thus, despite the emergence of resistance, continued CDK4/6i treatment markedly suppresses the growth rate of drug-resistant cells.

**Figure 1.**
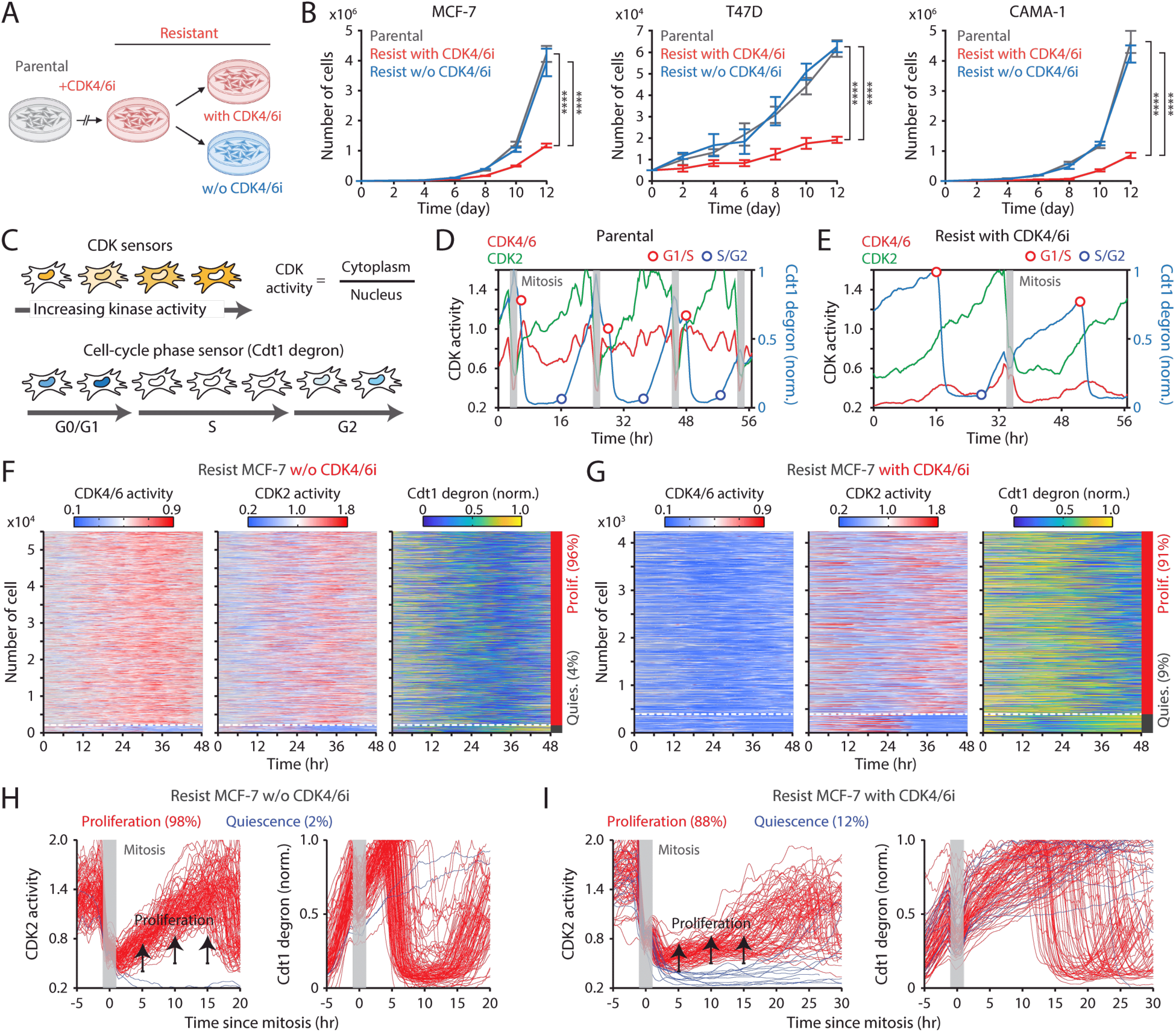
Continuous CDK4/6i treatment suppresses the growth of drug-resistant cells. (A) Schematic illustrating the establishment of CDK4/6i-resistant cells, either maintained in drug or withdrawn from treatment, for use in subsequent experiments. (B) Growth curves of parental and drug-resistant cells. Palbociclib (1 µM) was either withdrawn or maintained in drug-resistant cells. Data represent mean ± SD (*n* = 3 biological replicates). Statistical significance was determined by two-way ANOVA with Tukey’s post-hoc analysis (**** *P* < 0.0001). (C) Schematic of live-cell reporters for CDK4/6 and CDK2 activities (top) and cell-cycle phase (bottom). (D, E) Representative single-cell traces showing CDK4/6 and CDK2 activities together with Cdt1-degron intensity in parental (D) and drug-resistant (E) MCF-7 cells. Resistant cells were maintained in palbociclib (1 µM). (F, G) Heatmaps of single-cell traces showing CDK4/6 (left) and CDK2 (middle) activities, and Cdt1-degron intensity (right) in drug-resistant cells without (F) or with (G) continuous palbociclib (1 µM). Proliferating cells were identified by sustained CDK2 activity (>1 for >2 hr during 30–48 hr). (H, I) Single-cell traces of CDK2 activity (left) and Cdt1-degron intensity (right), aligned to the end of mitosis (anaphase) in drug-resistant cells without (H) or with (I) continuous palbociclib (1 µM). Based on CDK2 activity (black line), cells were classified as proliferating (red) or quiescent (blue).

To investigate how continued CDK4/6i treatment impedes the growth of drug-resistant cells, we monitored cell-cycle progression at the single-cell level. We established MCF-7 and MDA-MB-231 cells expressing live-cell sensors for CDK4/6 activity (35), CDK2 activity (36), and cell-cycle phase (Cdt1 degron) (37), as well as a nucleus marker (histone 2B) for individual cell tracking. The CDK sensors dynamically shuttle between the nucleus and cytoplasm depending on phosphorylation by their respective kinases (Figure 1C top). The Cdt1 degron is degraded during the S phase, allowing for the visualization of the G1/S and S/G2 transitions (Figure 1C, bottom). In parental cells, CDK4/6 was continuously activated after mitosis, followed by a gradual activation of CDK2, leading to S-phase entry (Figure 1D). In contrast, drug-resistant cells entered S phase through CDK2 activation with minimal CDK4/6 activity (Figure 1E). When evaluating thousands of cells, about 98% of parental cells exhibited sustained CDK4/6 and CDK2 activation to enter the cell cycle (Figure S1C and S1D). While drug withdrawal reactivated CDK4/6, over 90% of drug-resistant cells continued to proliferate regardless of the presence of CDK4/6i (Figure 1F, 1G, S1E, and S1F). To further assess cell-cycle dynamics, we aligned cells at cytokinesis and classified them as proliferating or quiescent based on CDK2 activity. In the absence of CDK4/6i, drug-resistant cells displayed CDK2/46 and CDK2 activity dynamics and S-phase entry kinetics comparable to those of parental cells (Figure 1H and S2A–S2D). By contrast, continued CDK4/6i treatment in resistant cells resulted in persistently low CDK4/6 activity, delayed CDK2 activation kinetics, and greater heterogeneity in the timing of S-phase entry, as indicated by Cdt1 degradation (Figure 1I, S2E, and S2F). These results demonstrate that despite active proliferation, continuous CDK4/6i treatment markedly decelerates and destabilizes cell-cycle progression in resistant cells, thereby limiting their growth rates.

CDK6 overexpression has emerged as a key driver of CDK4/6i resistance (38–41). In line with these findings, we observed elevated CDK6 levels in CDK4/6i-resistant cells (Figure S3A). To investigate the role of CDK6 in modulating CDK4/6 activity under resistance, we established CDK6 knockout (KO) MCF-7 cells using CRISPR-Cas9 and induced resistance to CDK4/6i (Figure S3B). Despite CDK6 depletion, CDK4/6 activity remained comparable between wild-type and CDK6-KO cells under continuous CDK4/6i treatment (Figure S3C–S3F). These results suggest a potential non-canonical role for CDK6 in mediating resistance to CDK4/6i therapy.

To assess the impact of continuous CDK4/6i treatment on cell-cycle progression, we measured the intermitotic time and the duration of each cell-cycle phase in MCF-7 and CAMA-1 cells (Figure 2A). The intermitotic time was comparable between parental and drug-resistant cells without ongoing CDK4/6i treatment (Figure 2B). However, in drug-resistant cells that were continuously treated with CDK4/6i, the intermitotic time was extended by 30–50% compared to other conditions. Moreover, continuous CDK4/6i treatment prolonged the G1 phase by approximately 200–300% without affecting the duration of the S and G2/M phases (Figure 2C–2E). Our data indicate that maintaining CDK4/6i treatment in drug-resistant cells significantly extends the G1 phase, thereby decelerating the overall growth rate.

**Figure 2.**
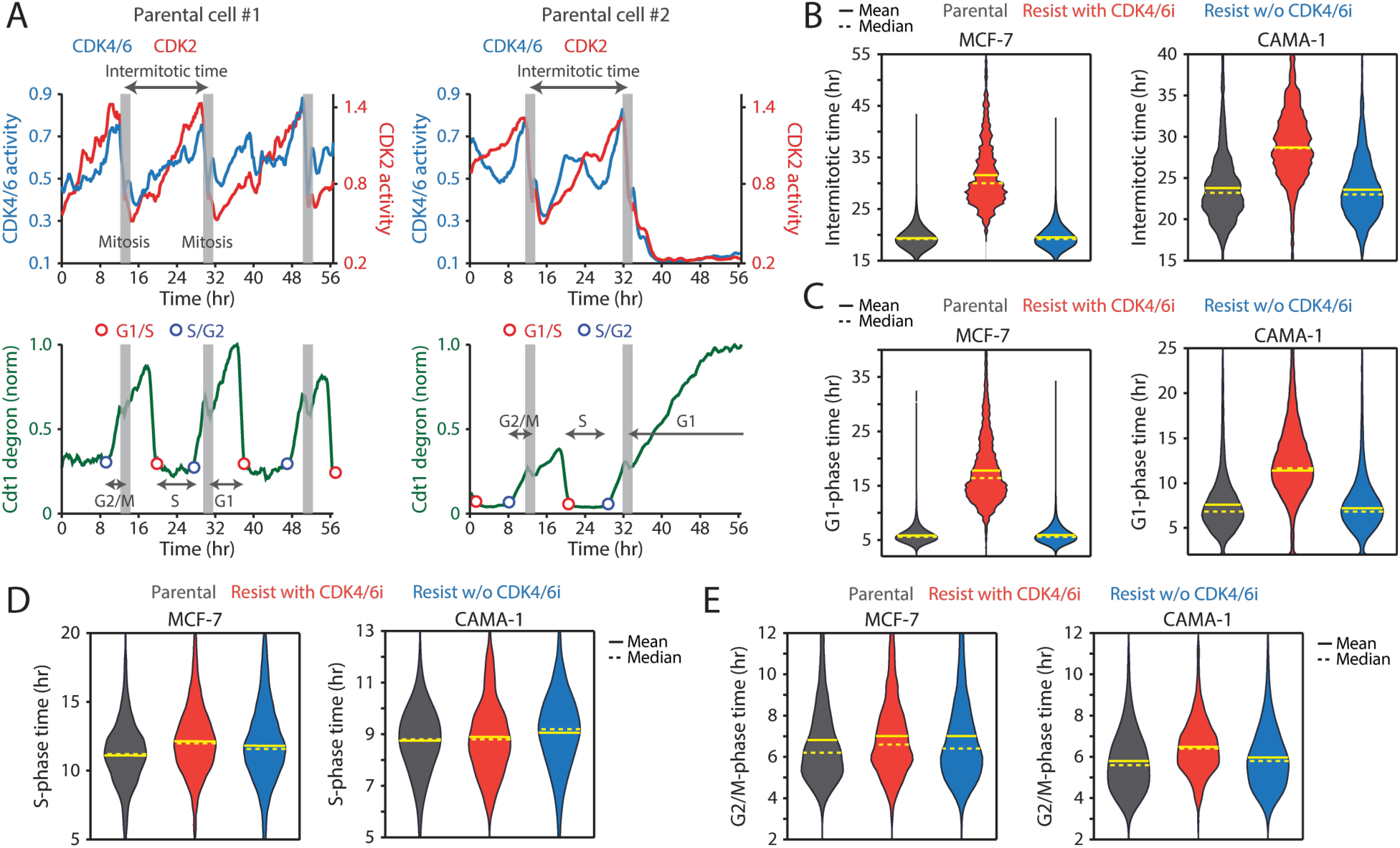
CDK4/6i maintenance extends G1-phase progression. (A) Representative single-cell traces showing CDK4/6 and CDK2 activities (top) and Cdt1-degron intensity (bottom) in parental MCF-7 cells. (B–E) Violin plots showing intermitotic time (*n* > 600 cells/condition) (B), G1-phase duration (*n* > 2,000 cells/condition) (C), S-phase duration (*n* > 400 cells/condition) (D), and G2/M-phase duration (*n* > 250 cells/condition) (E) in MCF-7 (left) and CAMA-1 (right) cells. Solid and dashed yellow lines indicate mean and median, respectively.

### Maintaining CDK4/6i treatment triggers ineffective Rb inactivation

We sought to elucidate the molecular mechanisms underlying the extended G1 phase and slow CDK2 activation kinetics in drug-resistant cells under continuous CDK4/6i treatment. Our recent studies demonstrated that CDK4/6 inhibition initially halts proliferation via Rb activation but ultimately reduces Rb protein due to decreased stability (42–44). Consistent with this, we found that drug-resistant cells maintained on CDK4/6i exhibited reduced levels of both total and phosphorylated Rb compared to parental cells (Figure 3A). Importantly, CDK4/6i withdrawal restored both total and phosphorylated Rb levels, confirming that the effect was reversible. However, Rb loss in resistant cells was incomplete, as demonstrated by comparisons with Rb-KO cells (Figure S4A). To directly test the role of Rb, we used Rb-KO cells resistant to CDK4/6i. Remarkably, Rb KO rescued the prolonged G1 duration and restored rapid CDK2 activation kinetics without affecting CDK4/6 activity, even under continuous CDK4/6 inhibition (Figures S4B and S4C). These findings led us to hypothesize that maintaining CDK4/6i induces a non-canonical pathway of Rb inactivation via post-translational degradation. This passive inactivation may result in inefficient E2F activation and delayed CDK2 engagement, thereby slowing G1-phase progression.

**Figure 3.**
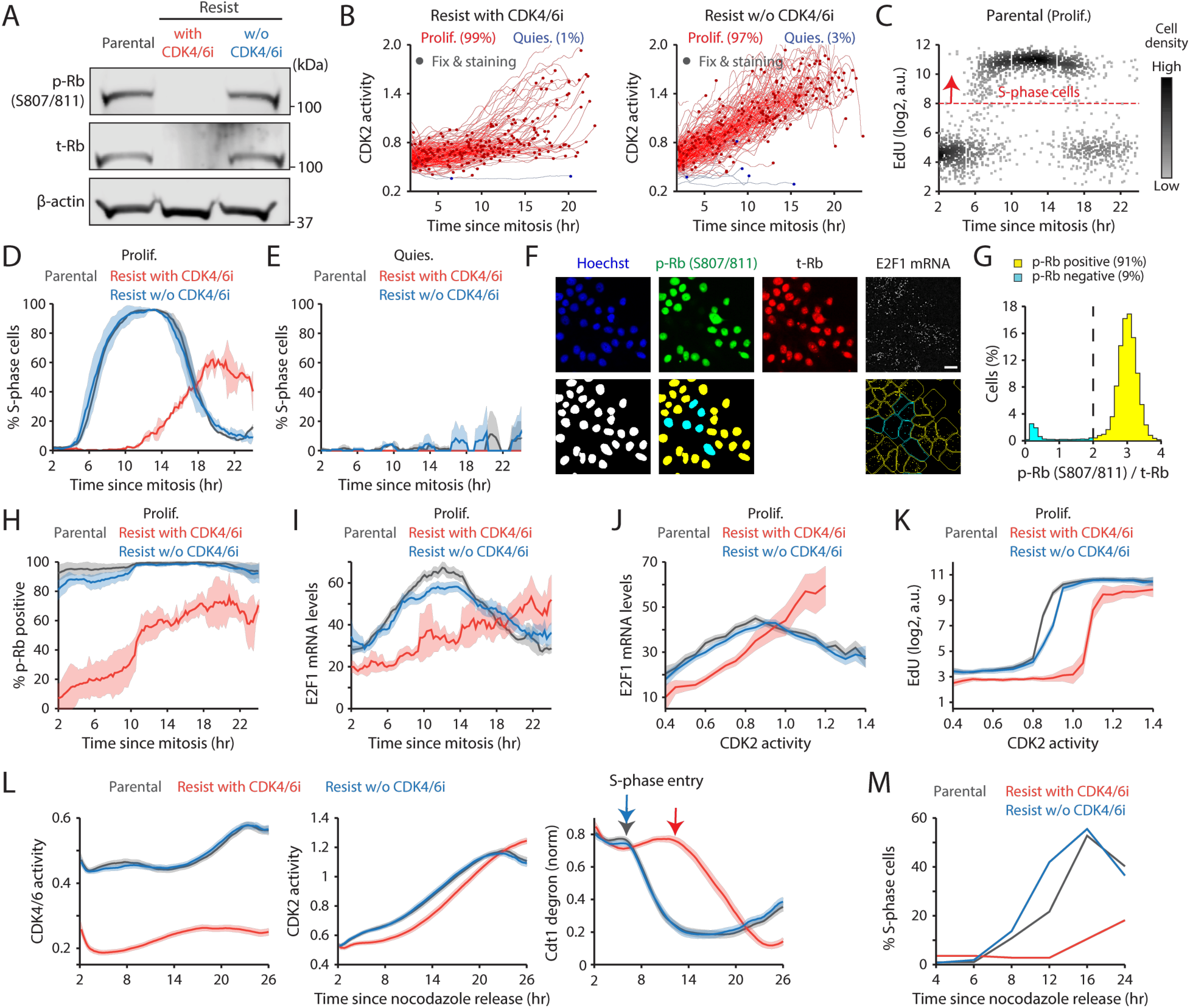
CDK4/6i maintenance induces an ineffective Rb inactivation pathway. (A) Immunoblot of phosphorylated Rb (S807/811; p-Rb), total Rb (t-Rb), and β-actin protein levels in MCF-7 cells. Drug-resistant cells were harvested two weeks after drug withdrawal. (B) Singe-cell traces of CDK2 activity aligned to mitosis in proliferating (red) and quiescent (blue) MCF-7 cells. Circles indicate the time of fixation and staining (*n* = 200 cells). (C) Scatter plot of EdU intensity versus time since mitosis in parental cells. The red dotted line marks the threshold for S phase (*n* = 2,000 cells). (D, E) Percentage of S-phase cells as a function of time since mitosis in proliferating (D) and quiescent (E) cells. Data represent mean ± SD (*n* = 2 biological replicates). (F) Representative immunostaining of Hoechst, p-Rb, t-Rb, and E2F1 mRNA FISH (top). Processed images showing nucleus segmentation, p-Rb classification, and mRNA puncta detection (bottom). (G) Histogram of cells with p-Rb normalized to t-Rb. (H, I) Percentage of cells with p-Rb (H) and E2F1 mRNA levels (I) as a function of time since mitosis. Data represent mean ± SD (H, *n* = 3 biological replicates) or mean ± 95% confidence intervals (I, *n* > 2,500 cells/condition). (J, K) E2F1 mRNA (J) and EdU (K) levels as a function of CDK2 activity. Data represent mean ± 95% confidence intervals (*n* > 2,500 cells/condition). (L, M) Averaged traces of CDK4/6 (left) and CDK2 (middle) activities and Cdt1-degron intensity (right) (L) and percentage of S-phase cells (M). MCF-7 cells were released after synchronizing with nocodazole (250 nM, 14 hr).

To test this hypothesis, we tracked CDK2 activity in individual cells and aligned proliferative events with fixed-cell measurements from the end of mitosis (Figure 3B and S5A). Using 5-ethynyl-2’-deoxyuridine (EdU) incorporation, we quantified the kinetics of S-phase entry. Parental and drug-resistant cells without CDK4/6i transitioned sharply into S phase approximately 6 hr after mitosis and completed DNA replication within 12 hr (Figures 3C, 3D, and S5B). In contrast, drug-resistant cells under sustained CDK4/6 inhibition exhibited markedly delayed and heterogeneous G1/S transitions (Figures 3D and S5C). Quiescent cells, as expected, showed no EdU incorporation (Figure 3E). We next assessed the dynamics of Rb phosphorylation and E2F activity using immunostaining and mRNA FISH (Figure 3F). Rb hyperphosphorylation was classified by phosphorylation at Ser807/811, a well-characterized marker (45) (Figure 3G). E2F1 mRNA levels, which are autoregulated (46), serve as a proxy for E2F transcriptional output (8,45,47). In the absence of CDK4/6i, both parental and drug-resistant cells robustly induced Rb phosphorylation before S-phase entry (Figure 3H). Conversely, under continuous CDK4/6 inhibition, resistant cells initiated CDK2 activation and G1 progression without Rb hyperphosphorylation, resulting in slower and weaker E2F1 mRNA induction (Figures 3H and 3I). When comparing CDK2 activity with E2F1 mRNA levels or EdU incorporation, resistant cells under CDK4/6i maintenance required higher CDK2 activity to reach equivalent E2F1 mRNA levels or S-phase entry (Figure 3J, 3K, S5D, and S5E). The reduced E2F1 mRNA levels during S phase likely result from suppression by atypical E2Fs, E2F7 and E2F8 (48). Finally, mitotic synchronization with nocodazole confirmed that CDK4/6i maintenance prolongs G1 and delays S-phase entry in resistant cells (Figure 3L and 3M). Together, these findings indicate that continuous CDK4/6 inhibition in drug-resistant cells triggers inefficient Rb inactivation, attenuated E2F induction, and delayed CDK2-driven G1/S transition, leading to prolonged and heterogeneous G1-phase progression.

### Maintenance of CDK4/6i suppresses the growth of drug-resistant tumors

We investigated the therapeutic benefits of maintaining CDK4/6i treatment after the onset of drug resistance in vivo. To this end, we established an MCF-7 xenograft model by orthotopically inoculating MCF-7 cells into the mammary fat pad of immunodeficient mice. Palbociclib was initially administered when tumors reached 100 mm^3^ (Figure 4A). Following an initial period of tumor stasis/regression of variable duration, tumors resumed growth, indicating the development of acquired resistance (Figure 4B). Upon regrowth to 145–155 mm^3^, mice were randomized to one of four secondary regimens: treatment discontinuation, continued palbociclib, or a switch to ribociclib or abemaciclib. To mirror clinical practice, ribociclib was administered at a fourfold higher dose than the other CDK4/6i. Consistent with our in vitro findings, continuation of any CDK4/6i significantly attenuated overall tumor growth relative to treatment discontinuation (Figure 4C–E). Additionally, we observed that switching to ribociclib or abemaciclib resulted in significantly greater tumor control compared to continuing palbociclib. These observations align with clinical studies reporting agent-specific differences in second-line PFS (19–21,23). Notably, analysis of tumor tissues after secondary treatment showed restoration of total Rb expression only in the drug-discontinued cohort (Figure 4F). Together, these data support maintaining CDK4/6 inhibition to restrain the overall growth of drug-resistant tumors.

**Figure 4.**
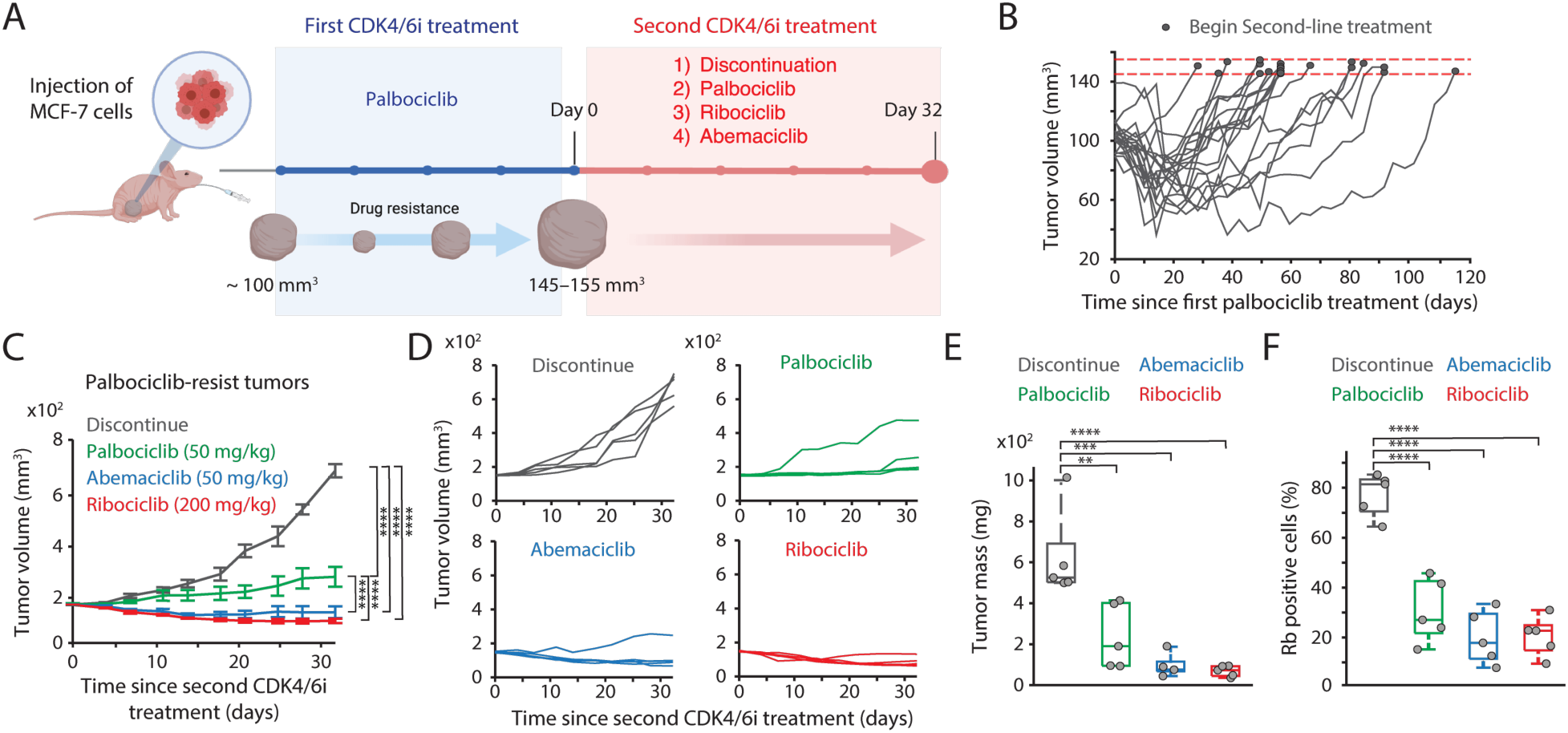
CDK4/6i maintenance suppresses the growth of drug-resistant tumors. (A) Schematic representation of experimental design. Once tumors reached a volume of 100 mm^3^, mice were treated with palbociclib. Following the development of resistance, mice were randomly assigned to one of four treatment groups: treatment discontinuation, palbociclib maintenance, switch to ribociclib, or switch to abemaciclib for 32 days. (B) Tumor growth curves showing the establishment of resistance to palbociclib. Horizontal red dotted lines (145−155 mm^3^) indicate the point at which mice were assigned to second CDK4/6i treatment (*n* = 20 mice). (C, D) Averaged (C) and individual (D) tumor growth traces following second CDK4/6i treatment. Data represent mean ± SEM (*n* = 5 mice/condition). Statistical significance was determined using two-way ANOVA with Tukey’s post-hoc analysis (**** *P* < 0.0001). (E, F) Box plots showing tumor mass (E) and the percentage of Rb-positive cells (F) after second CDK4/6i treatment (*n* = 5 mice/condition). Statistical significance was determined using one-way ANOVA with Tukey’s post-hoc analysis (** *P* < 0.05, *** *P* < 0.01, **** *P* < 0.0001).

### Addition of ET augments the efficacy of CDK4/6i maintenance

Our recent studies have highlighted the pivotal role of c-Myc in amplifying E2F activity and fostering CDK4/6i resistance following alternative Rb inactivation (42,43). Using a doxycycline-inducible system, we found that induction of c-Myc significantly facilitated the growth of drug-resistant cells under ongoing CDK4/6i treatment (Figure S6A). Furthermore, despite similar low CDK4/6 activity, c-Myc induction facilitated cell-cycle progression in CDK4/6i-resistant cells (Figure S6B–D). These data demonstrated that c-Myc promotes CDK4/6i resistance by facilitating cell-cycle progression in drug-resistant cells.

Given the estrogen responsiveness of c-Myc (49), we hypothesized that the continued addition of ET could further suppress the growth of CDK4/6i-resistant cells by downregulating c-Myc expression. We employed the estrogen receptor antagonist fulvestrant to evaluate the therapeutic benefit of maintaining the combination of CDK4/6i and ET. While MCF-7 cells exhibited primary resistance to fulvestrant, CAMA-1 and T47D cells displayed sensitivity to this drug (Figure 5A). We used these MCF-7 and CAMA-1 cells to examine whether primary ET resistance contributes to the benefit of maintaining CDK4/6i and ET in drug-resistant cells. To establish drug resistance, CDK4/6i-resistant cell lines were chronically treated with a combination of palbociclib and fulvestrant for over two months. Subsequently, these resistant cell lines were subjected to continuous treatment with either palbociclib or fulvestrant alone or in combination, and their cumulative proliferation rate was monitored. We observed a significantly slower growth rate in drug-resistant cells continuously exposed to the combination therapy compared to other treatment conditions, not only in CAMA-1 but also in MCF-7 cells (Figure 5B, S7A, and S7B). Continued treatment with fulvestrant, either alone or combined with palbociclib, reduced c-Myc levels compared to palbociclib maintenance alone (Figure 5C). Our data indicate distinct resistance mechanisms between CDK4/6i and ET. Therefore, regardless of primary resistance to ET, the continued addition of ET to CDK4/6i treatment is beneficial in suppressing the growth of drug-resistant cells by inhibiting c-Myc.

**Figure 5.**
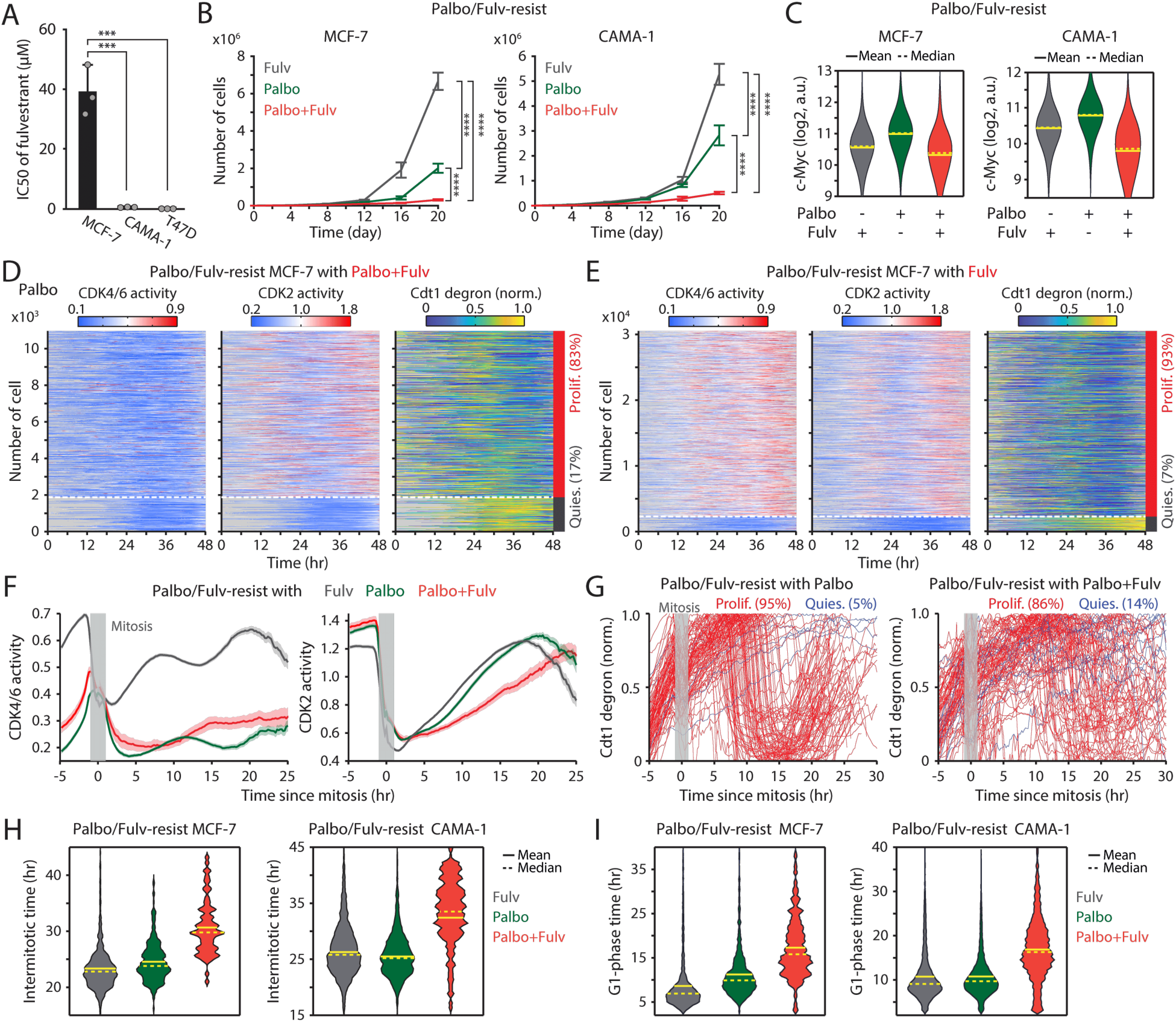
Maintaining CDK4/6i and ET synergistically suppresses the growth of drug-resistant cells. (A) IC50 values of fulvestrant. Data represent mean ± SD (*n* = 3 biological replicates). Statistical significance was determined by one-way ANOVA with Tukey’s post-hoc analysis (*** *P* < 0.001). Growth curves of MCF-7 (left) and CAMA-1 (right) cells resistant to palbociclib and fulvestrant. Cells were maintained with fulvestrant (500 nM), palbociclib (1 µM), or the combination. Data represent mean ± SD (*n* = 3 biological replicates). Statistical significance was determined by two-way ANOVA with Tukey’s post-hoc analysis (**** *P* < 0.0001). (C) Violin plots of c-Myc levels measured by immunofluorescence in resistant cells resistant treated with the indicated drugs for one week. Solid and dashed yellow lines represent mean and median, respectively (*n* > 2,000 cells/condition). (D, E) Heatmaps showing single-cell traces for CDK4/6 (left) and CDK2 (middle) activities and Cdt1-degron intensity (right) in drug-resistant cells maintained with palbociclib (1 µM) and fulvestrant (500 nM) (D) or fulvestrant alone (E). Proliferating cells were identified based on CDK2 activity (>1 for more than 2 hr between 30 and 48 hr). (F) Averaged traces of CDK4/6 (left) and CDK2 (right) activities aligned by mitosis in drug-resistant cells treated with the indicated drugs. Data represent mean ± 95% confidence intervals (*n* > 1,000 cells/condition). (G) Single-cell traces of Cdt1-degron intensity aligned by mitosis in drug-resistant cells treated with palbociclib (1 µM) (left) or palbociclib + fulvestrant (500 nM) (right). (H, I) Violin plots showing intermitotic time (*n* > 70 cells/condition) (H), and G1-phase duration (*n* > 630 cells/condition) (I) in MCF-7 (left) and CAMA-1 (right) cells. Solid and dashed yellow lines indicate mean and median, respectively.

To elucidate the mechanisms underlying the attenuation of growth resulting from continued treatment with the drug combination, we assessed CDK4/6 and CDK2 activities along with cell-cycle progression (Cdt1 degron) in drug-resistant MCF-7 cells. Cells were continuously treated with palbociclib or fulvestrant alone or in combination. By classifying cells into proliferation and quiescence based on CDK2 activation, we found over 80% of cells were proliferating in all conditions (Figure 5D, 5E, and S7C). Furthermore, we observed reactivation of CDK4/6 in drug-resistant cells continuously treated with fulvestrant alone. However, when we monitored cell-cycle dynamics by aligning cells to mitosis, the maintenance of the drug combination caused slower CDK2 activation kinetics and greater heterogeneity in S-phase entry, as indicated by Cdt1 degradation, compared to single-drug treatment conditions (Figure 5F, 5G, S7D, and S7E). By analyzing each cell-cycle duration, we observed a significant increase in intermitotic time and G1-phase duration, while S- or G2-phase lengths remained relatively unchanged (Figure 5H, 5I, S7F, and S7G). This emphasizes the role of the drug combination in altering cell-cycle kinetics in the G1 phase. Our findings showed the therapeutic benefits of maintaining a combination of CDK4/6i and ET in attenuating the growth of drug-resistant cells.

### The benefit of combining CDK2i with CDK4/6i as a second-line therapy

We next evaluated the therapeutic potential of adding the CDK2i INX-315 as a second-line strategy. Palbociclib/fulvestrant-resistant MCF-7 and CAMA-1 cells were treated under five regimens: treatment discontinuation, fulvestrant alone, fulvestrant + palbociclib, fulvestrant + INX-315, or fulvestrant + palbociclib + INX-315. Continuous palbociclib + fulvestrant treatment significantly reduced resistant-cell growth compared to discontinuation, fulvestrant alone, or fulvestrant + INX-315 (Figure 6A–6C). Notably, the triple combination provided the most robust suppression of growth. To dissect cell-cycle dynamics, we assessed CDK activities and Cdt1 degron levels one week after treatment. Consistent with earlier results, palbociclib withdrawal triggered CDK4/6 reactivation (Figure S8A–S8D). Furthermore, we found CDK2 reactivation even in the presence of INX-315 (Figure 6D and S8D). While CDK2 reactivation was also observed, the triple combination produced the longest intermitotic intervals and G1-phase duration without affecting the S and G2 phases (Figure 6E and S8E). Analysis of cells synchronized by mitosis further confirmed that the triple-drug regimen most strongly delayed CDK2 activation kinetics (Figure 6F). Moreover, dose titration studies revealed a synergistic interaction between CDK4/6i and CDK2i (Figure 6G and S8F). To explore transcriptomic consequences, we performed RNA sequencing 20 days post-treatment. Continued palbociclib + fulvestrant suppressed E2F target genes more effectively than INX-315, and the triple combination achieved the most potent inhibition of both E2F and Myc target gene programs (Figure 6H and S8G). Together, these findings demonstrate that, despite partial CDK2 reactivation, adding CDK2i to ongoing CDK4/6i + ET further attenuates E2F activity, delays CDK2 engagement, and suppresses resistant-cell growth. These results support a therapeutic rationale for maintaining CDK4/6 blockade while strategically incorporating CDK2 inhibition after progression.

**Figure 6.**
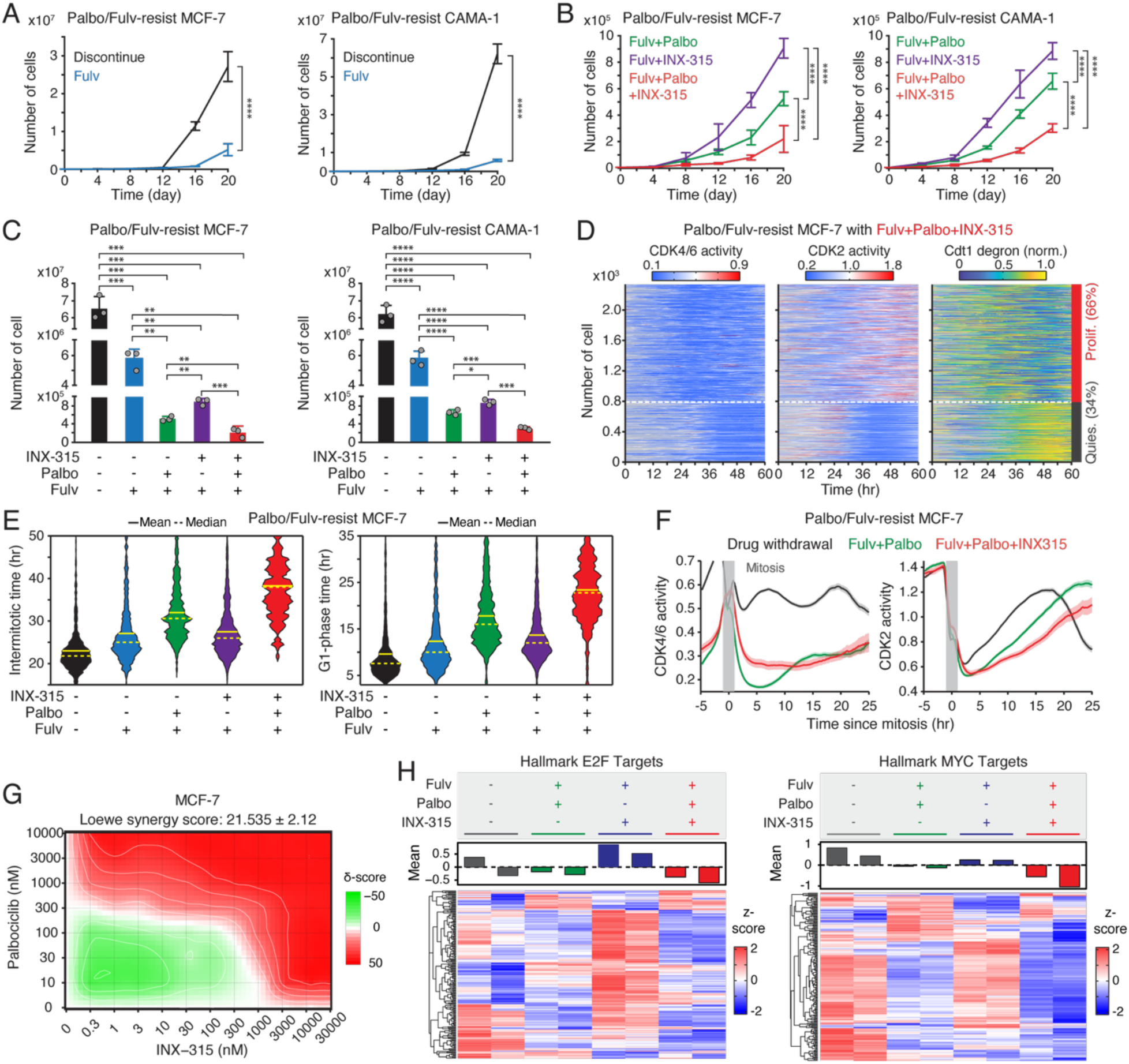
Therapeutic benefit of combining CDK2i with CDK4/6i and ET. (A, B) Growth curves of MCF-7 (left) and CAMA-1 (right) cells resistant to palbociclib and fulvestrant under various treatment conditions: drug discontinuation, fulvestrant (500 nM) alone, or in combination with palbociclib (1 µM) and/or INX-315 (100 nM). Data represent mean ± SD (*n* = 3 biological replicates). *P* values were calculated with two-way ANOVA with Tukey’s post-hoc analysis (**** *P* < 0.0001). (C) Cell numbers 20 days after drug treatment. Data represent mean ± SD (*n* = 3 biological replicates). *P* values were calculated using an unpaired *t*-test (* *P* < 0.05, ** *P* < 0.01, *** *P* < 0.001, **** *P* < 0.0001). (D) Heatmaps of single-cell traces for CDK4/6 (left) and CDK2 (middle) activities, and Cdt1-degron intensity (right) in palbociclib/fulvestrant resistant MCF-7 cells treated with the triple combination of palbociclib (1 µM), fulvestrant (500 nM), and INX-315 (100 nM) for one week before imaging. Proliferating cells were identified based on CDK2 activity (>1 for more than 2 hr between 30 and 48 hr). (E) Violin plots showing intermitotic time (*n >* 200 cells/condition) (left) and G1-phase duration (*n >* 900 cells/condition) (right). Solid and dashed yellow lines indicate mean and median, respectively. (F) Averaged traces of CDK4/6 (left) and CDK2 (right) activities aligned by mitosis in drug-resistant cells treated with the indicated drugs. Data represent mean ± 95% confidence intervals (*n* > 1,000 cells/condition). (G) Loewe synergy score calculated from percent-inhibition data generated by dual titration of palbociclib (0–10µM) and INX-315 (0–30µM) in MCF-7 cells following 48 hr treatment (*n* = 3 biological replicates). (H) Heatmaps comparing gene expression profiles for hallmark E2F (left) and MYC (right) targets in drug-resistant MCF-7 cells treated with the indicated drugs for 20 days. Samples were collected as biological duplicates.

### Role of cyclin E/A in CDK2 reactivation under combined CDK4/6 and CDK2 inhibition

A recent study identifies the cyclin A-CDK complex as a key driver of rapid adaptation to CDK2 inhibition (32). To investigate the mechanism of CDK2 reactivation during CDK2i treatment, we explored the role of the CDK2 activators, cyclins E and A. Using a doxycycline-inducible system, we induced the overexpression of cyclin E1 or A2 in drug-naïve MCF-7 cells (Figure S9A). While treatment with INX-315 (100 nM–1 µM) initially inhibited CDK2 activity, CDK2 reactivation occurred within 1–2 hr after drug treatment, regardless of cyclin E1 or A2 overexpression (Figure 7A and S9B–S9D). The combination of CDK2i and CDK4/6i blocked this rapid CDK2 reactivation, but the overexpression of cyclin E1 or A2 hindered complete suppression (Figure 7B and S9E–S9G). These findings suggest that CDK4/6 inhibition is necessary for effectively suppressing CDK2 activity by CDK2i, and that cyclin E1 and A2 overexpression may contribute to CDK2 reactivation.

**Figure 7.**
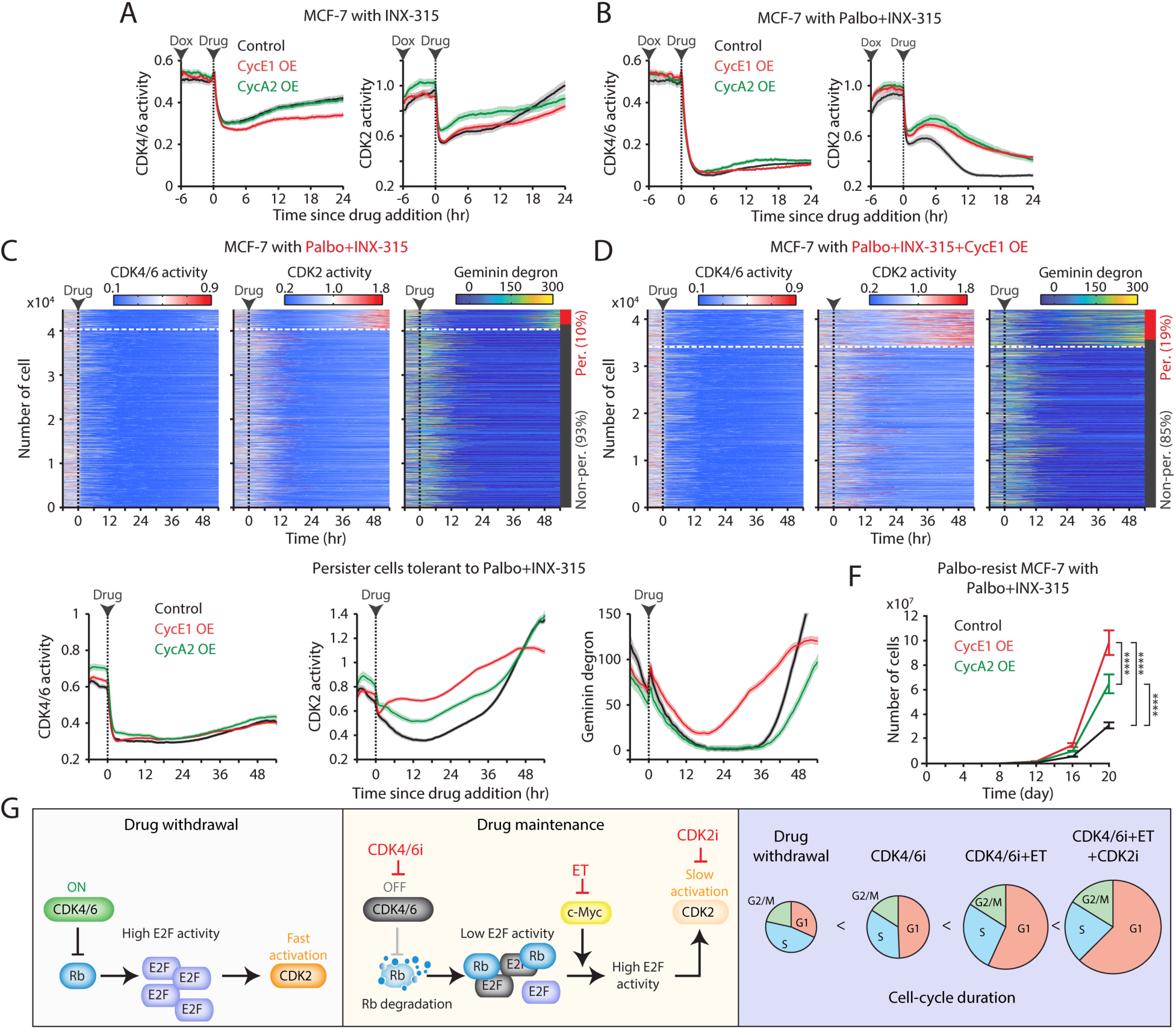
Overexpression of cyclin E and A facilitates resistance to the combination of CDK2i and CDK4/6i. (A, B) Averaged traces of CDK4/6 (left) and CDK2 (right) activities in MCF-7 cells with or without cyclin E1 or A2 overexpression. Cells were treated with doxycycline (500 nM) 6 hr before the addition of INX-315 (1 µM) alone (A) or in combination with palbociclib (1 µM) (B). Data represent mean ± 95% confidence intervals (*n* > 900 cells/condition). (C, D) Heatmaps of single-cell traces for CDK4/6 (left) and CDK2 (middle) activities, and Geminin-degron intensity (right) in drug-naïve MCF-7 cells without (C) or with (D) cyclin E1 overexpression. Cells were treated with palbociclib (1 µM), INX-315 (100 nM), and doxycycline (500 nM). Persister cells were identified based on CDK2 activity (>1 for more than 2 hr between 30 and 48 hr). (E) Averaged traces of CDK4/6 (left) and CDK2 (middle) activities and Geminin degron intensity (right) in persister cells tolerant to combination palbociclib (1 µM), and INX-315 (100 nM) without or with cyclin E1 or A2 overexpression. Data represent mean ± 95% confidence interval. *n* > 2500 cells/condition. (F) Growth curves of palbociclib-resistant MCF-7 cells without or with cyclin E1 or A2 overexpression. Cells were treated with palbociclib (1 µM), INX-315 (100 nM), and doxycycline (500 nM). Data represent mean ± SD (*n* = 3 biological replicates). Statistical significance was determined using two-way ANOVA with Tukey’s post-hoc analysis (**** *P* < 0.0001). (G) Summary schematic illustrating the mechanisms underlying the benefits of continued CDK4/6i and ET therapies and the introduction of CDK2i in drug-resistant cells.

We next examined the impact of cyclin E and A overexpression in the development of persister cells, which are implicated in residual tumor growth and eventual resistance (42,50). We used drug-naïve MCF-7 cells expressing live-cell sensors for CDK2, CDK4/6, and anaphase-promoting complex/cyclosome (APC/C) (Geminin degron) activities (51). APC/C is a multi-subunit E3 ubiquitin ligase typically inactivated near the G1/S transition, leading to Geminin-degron accumulation. Treatment with CDK4/6i and CDK2i induced near-complete cell-cycle arrest within 24 hr (Figure 7C). However, approximately 10% of cells developed a persister phenotype through CDK2 reactivation. Overexpression of cyclin E1 increased the percentage of persister cells, while cyclin A2 had no obvious impact (Figure 7D and S10). Further kinetic analysis revealed that cyclin E1 overexpression accelerated CDK2 reactivation and APC/C inactivation in persister cells (Figure 7E). These findings indicate that cyclin E1, but not cyclin A2, facilitates CDK2 reactivation and persister development, promoting resistance to the CDK4/6i and CDK2i combination.

Since cyclin A is a substrate of APC/C (52), the lack of effect from cyclin A2 induction could be due to its degradation by APC/C. Thus, this degradation may reduce cyclin A to insufficient levels, preventing its impact on persister cell development. To further evaluate the impact of cyclin E and A overexpression on resistance development, we exposed palbociclib-resistant MCF-7 cells to the combination of palbociclib and INX-315. Both cyclin E1 and A2 overexpression significantly accelerated the development of resistance to the drug combination, with cyclin E1 having a more significant effect than cyclin A2 (Figure 7F). These results indicate that although CDK2 activity is targeted by CDK2i, the overexpression of cyclin E and A promotes resistance to the combination of CDK4/6i and CDK2i. Due to cyclin A degradation by APC/C in quiescent cells, cyclin E may play a more critical role in driving resistance to this drug combination.

## Discussion

The combination of CDK4/6i and ET has reshaped treatment for HR^+^/HER2^-^ breast cancer (53–60). However, resistance commonly emerges, and no consensus second-line standard is established. Our data show that continued CDK4/6i treatment in drug-resistant cells engages a non-canonical, proteolysis-driven route of Rb inactivation, yielding attenuated E2F output and a pronounced delay in G1 progression (Figure 7G). Concurrent ET further deepens this blockade by suppressing c-Myc-mediated E2F amplification, thereby prolonging G1 and slowing population growth. Importantly, CDK2 inhibition alone was insufficient to control resistant cells. Robust suppression of both CDK2 activity and resistant-cell growth required CDK2i in combination with CDK4/6i, consistent with prior reports supporting dual CDK targeting (27–34). Moreover, cyclin E blunted the efficacy of the CDK4/6i + CDK2i combination by reactivating CDK2. Together, these findings provide a mechanistic rationale for maintaining CDK4/6i beyond progression and support testing the combination of CDK4/6i and CDK2i, as evidenced by concordant in vitro and in vivo results.

Our data indicate that maintaining both CDK4/6i and ET synergistically decelerates cell-cycle progression in drug-resistant cells by further delaying CDK2 activation kinetics and the G1/S transition without affecting the S and G2 phases. This dual effect stems from CDK4/6i causing suboptimal Rb inactivation while ET suppresses the global transcription amplifier c-Myc, collectively leading to diminished E2F transcriptional activity. As a result, this reduced E2F activity lowers the expression of critical cell-cycle genes, such as cyclin E and A, extending the time needed for CDK2 activation. Given that CDK2 plays an essential role in initiating and advancing DNA replication (61,62), its delayed activation significantly prolongs the G1/S transition. Moreover, CDK2 activation also contributes to Rb phosphorylation and inactivation. High CDK2 activity is required to phosphorylate Rb, and CDK2-mediated Rb phosphorylation is tightly coupled with DNA replication timing (8,45). Thus, upon Rb phosphorylation by CDK2 at the G1/S transition, drug-resistant cells may effectively proceed through the cell cycle even under continued CDK4/6i treatment.

Clinical trials evaluating the efficacy of sustained CDK4/6i therapy predominantly use PFS as the primary endpoint (19–23). However, our findings suggest that drug-resistant tumors continue to proliferate despite CDK4/6i maintenance. Consequently, maintaining CDK4/6i appears to slow tumor growth rather than completely arrest it. This underscores the need for clinical trials to consider overall survival and tumor progression rates as more appropriate endpoints for assessing the true benefits of sustained CDK4/6i therapy. Furthermore, the distinct polypharmacology profiles among CDK4/6i (63), with ribociclib being the most specific and abemaciclib the least, may explain the varying therapeutic outcomes observed among these inhibitors (23,64).

Maintaining CDK4/6i treatment beyond progression may be particularly beneficial for about 70% of patients who do not acquire new genetic mutations (65). However, it is important to recognize that resistance to CDK4/6i often arises from mutations in genes associated with mitogenic or hormone-signaling pathways (65–69). These include mutations in *PIK3CA*, *ESR1*, *FGFR1*–*3*, and *HER2*, which have been linked to increased c-Myc expression (49,70,71). Additionally, previous studies have identified *FAT1* mutations as a driver of CDK4/6i resistance (39,72). These resistance mutations may reduce the efficacy of maintaining CDK4/6i and ET therapy. Moreover, about 4.7% of HR^+^/HER2^-^ breast cancer patients exhibit *Rb* mutations (65,67), making CDK4/6i treatment unlikely to be effective, thus making its continuation inadvisable in these cases.

In conclusion, our study provides mechanistic rationale for maintaining CDK4/6i together with ET after disease progression in HR^+^/HER2^-^ breast cancers that retain an intact Rb/E2F pathway. The combination of CDK4/6i and CDK2i can further provide durable growth suppression, consistent with prior studies (27–34). However, it is essential to acknowledge that CDK2/4/6 inhibition may promote whole-genome duplication (73), potentially fueling more aggressive tumor evolution. Finally, we identify cyclin E overexpression as a key driver of resistance to dual CDK4/6i and CDK2i therapy, providing a basis for biomarker-guided patient selection and the development of strategies to overcome therapeutic resistance.

---

**Supplementary Figure 1.**
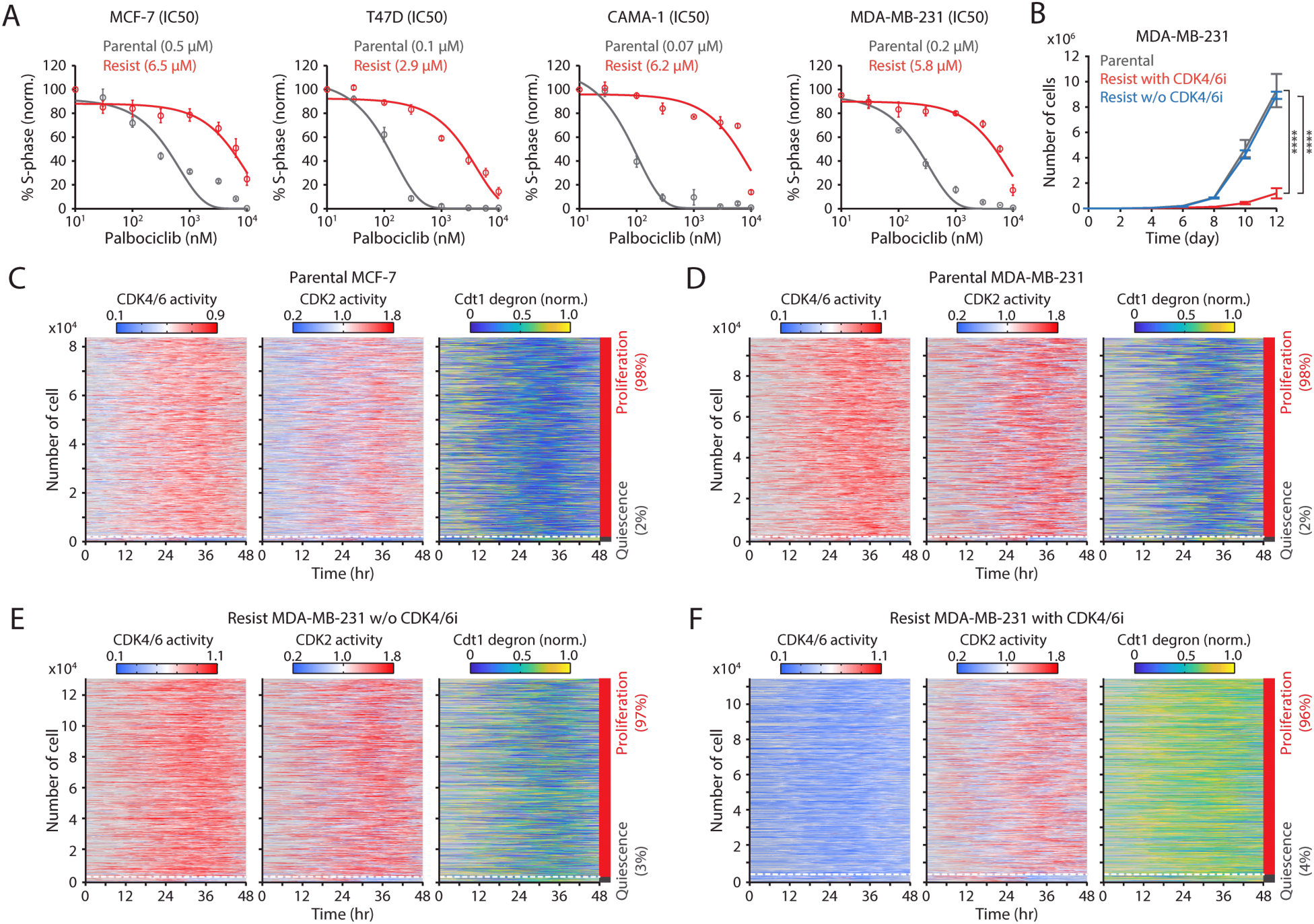
Validation of drug resistance and visualization of cell-cycle progression. (A) Dose-response curves of palbociclib showing its effects on the percentage of S-phase cells. Data represent mean ± SEM (*n* = 3 biological replicates). Solid lines represent sigmoidal best-fit curves. (B) Growth curves of drug-naïve and drug-resistant cells. Palbociclib (1 µM) was either withdrawn or maintained in drug-resistant cells. Data represent mean ± SD (*n* = 3 biological replicates). Statistical significance was determined using two-way ANOVA with Tukey’s post-hoc analysis (**** *P* < 0.0001). (C–F) Heatmaps of single-cell traces for CDK4/6 (left) and CDK2 (middle) activities, and Cdt1-degron intensity (right) in various conditions: drug-naïve MCF-7 (C) and MDA-MB-231 (D) cells, and drug-resistant MDA-MB-231 cells without (E) or with (F) continuous palbociclib (1µM) treatment. Proliferating cells were identified based on CDK2 activity (>1 for more than 2 hr between 30 and 48 hr).

**Supplementary Figure 2.**
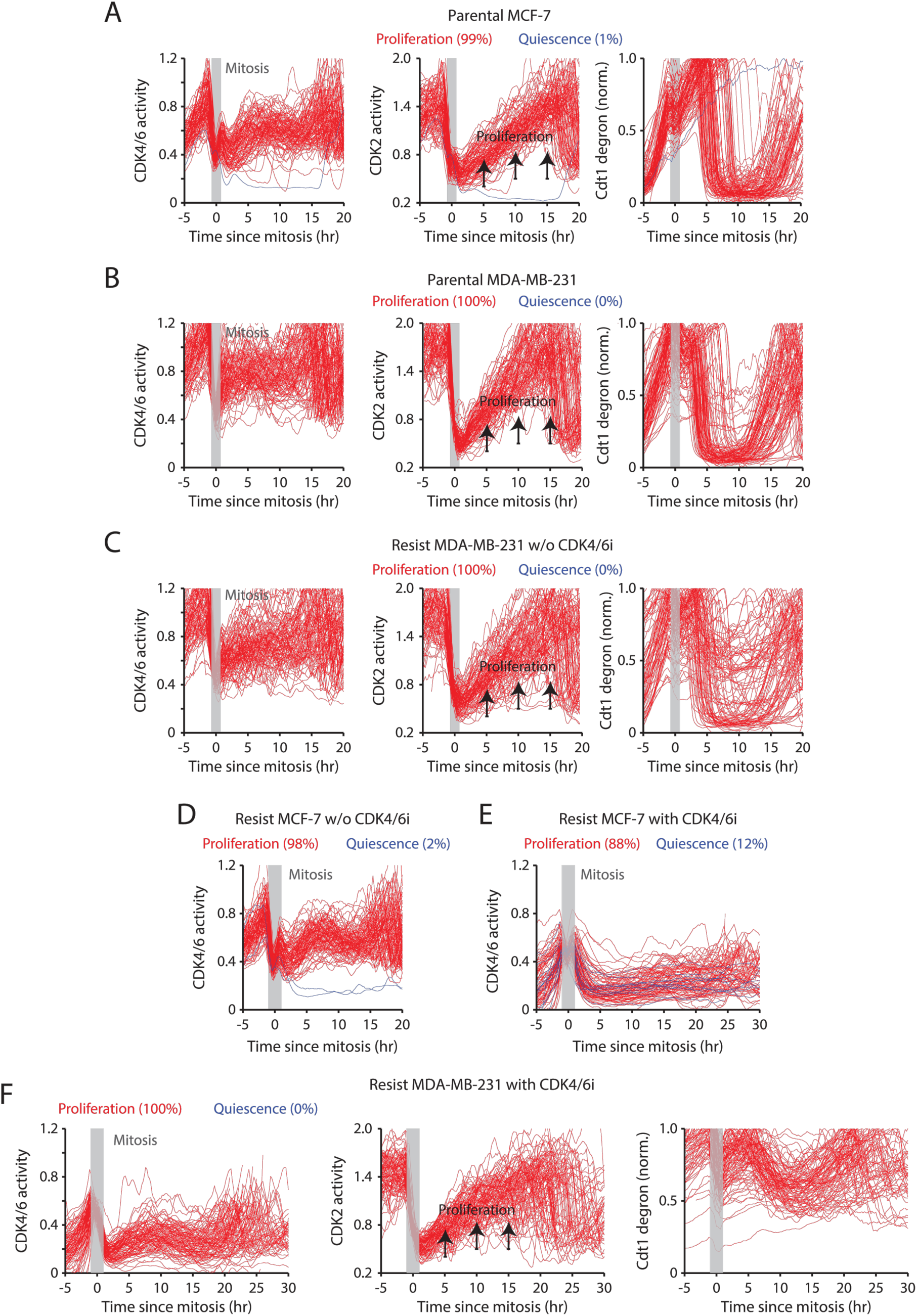
Slow cell-cycle progression in drug-resistant cells continuously treated with CDK4/6i. (A–C) Single-cell traces of CDK4/6 (left) and CDK2 (middle) activities and Cdt1-degron intensity (right) aligned by mitosis in drug-naïve MCF-7 (A) and MDA-MB-231 (B) cells and drug-resistant MDA-MB-231 cells without CDK4/6i treatment (C). The time of mitosis is marked in gray. (D, E) Single-cell trace of CDK4/6 activity corresponding to Figure 1G (D) and 1H (E). (F) Single-cell traces of CDK4/6 (left) and CDK2 (middle) activities and Cdt1-degron intensity (right) aligned by mitosis in drug-resistant MDA-MB-231 cells with continuous palbociclib (1 µM) treatment.

**Supplementary Figure 3.**
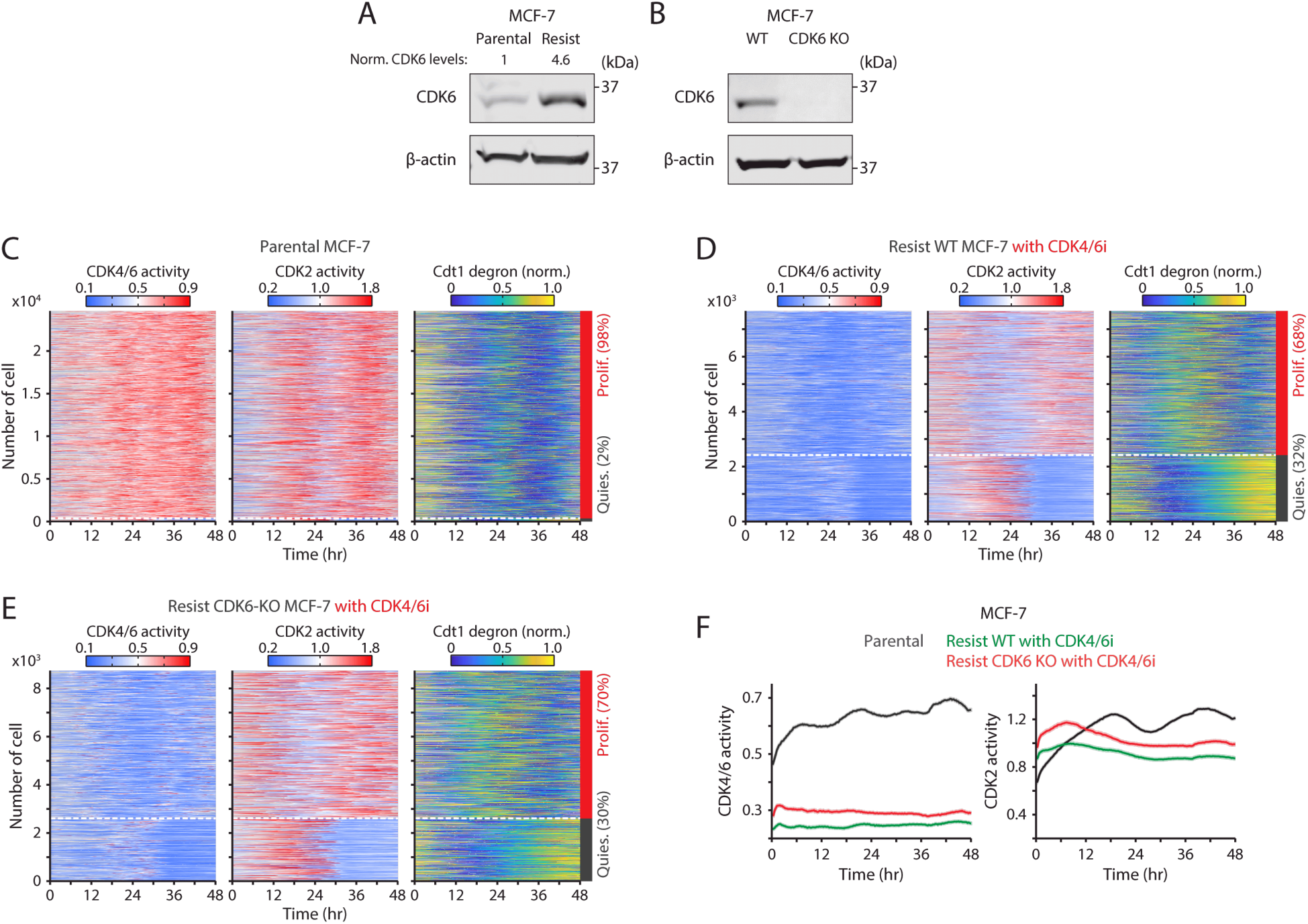
Potential non-canonical role of CDK6 in promoting CDK4/6i resistance. (A, B) Immunoblot showing CDK6 and β-actin expression in drug-naïve and palbociclib-resistant cells (A) and WT and CDK6-KO cells (B). The numerical values represent the intensity of the CDK6 band, normalized against the intensity of the β-actin band. (C–E) Heatmaps of single-cell traces for CDK4/6 (left) and CDK2 (middle) activities, and Cdt1 degron intensity (right) in drug-naïve (C) and drug-resistant WT (D) and CDK6-KO (E) cells with continued palbociclib (1 µM) treatment. Proliferating cells were identified based on CDK2 activity (>1 for >2 hr during 30–48 hr). (F) Averaged traces of CDK4/6 (left) and CDK2 (right) activities. Data represent mean ± 95% confidence intervals (*n* > 7,000 cells/condition).

**Supplementary Figure 4.**
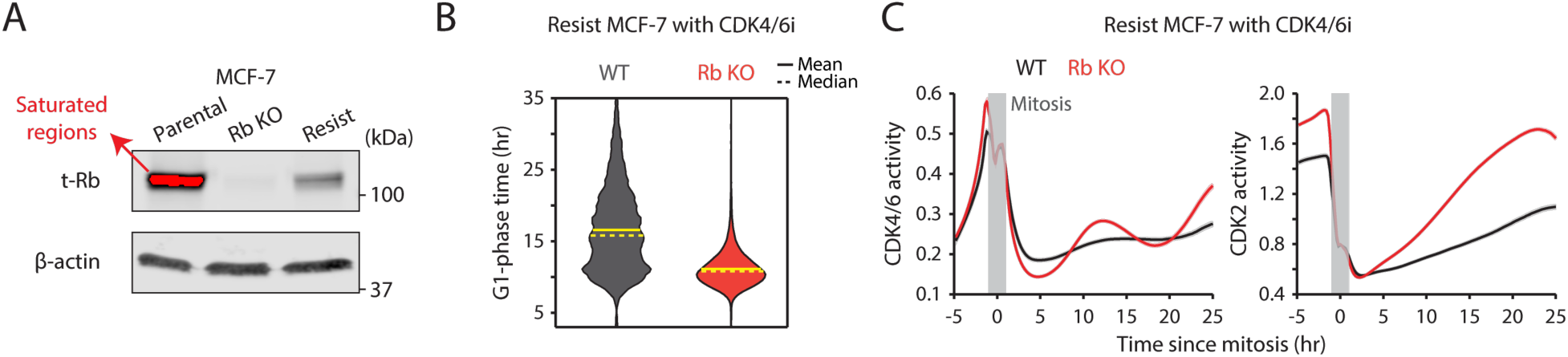
Incomplete Rb loss mediates the extended G1-phase progression. (A) Immunoblot showing total Rb and β-actin in WT, Rb-KO, and palbociclib-resistant cells. (B) G1-phase duration in drug-resistant WT and Rb-KO cells with continued palbociclib (1 µM) treatment. Solid and dashed yellow lines represent mean and median, respectively (*n >* 4,500 cells). (C) Averaged CDK4/6 (left) and CDK2 (right) activities aligned by mitosis in drug-resistant WT and Rb-KO cells with continued palbociclib (1 µM) treatment (*n >* 7,500 cells).

**Supplementary Figure 5.**
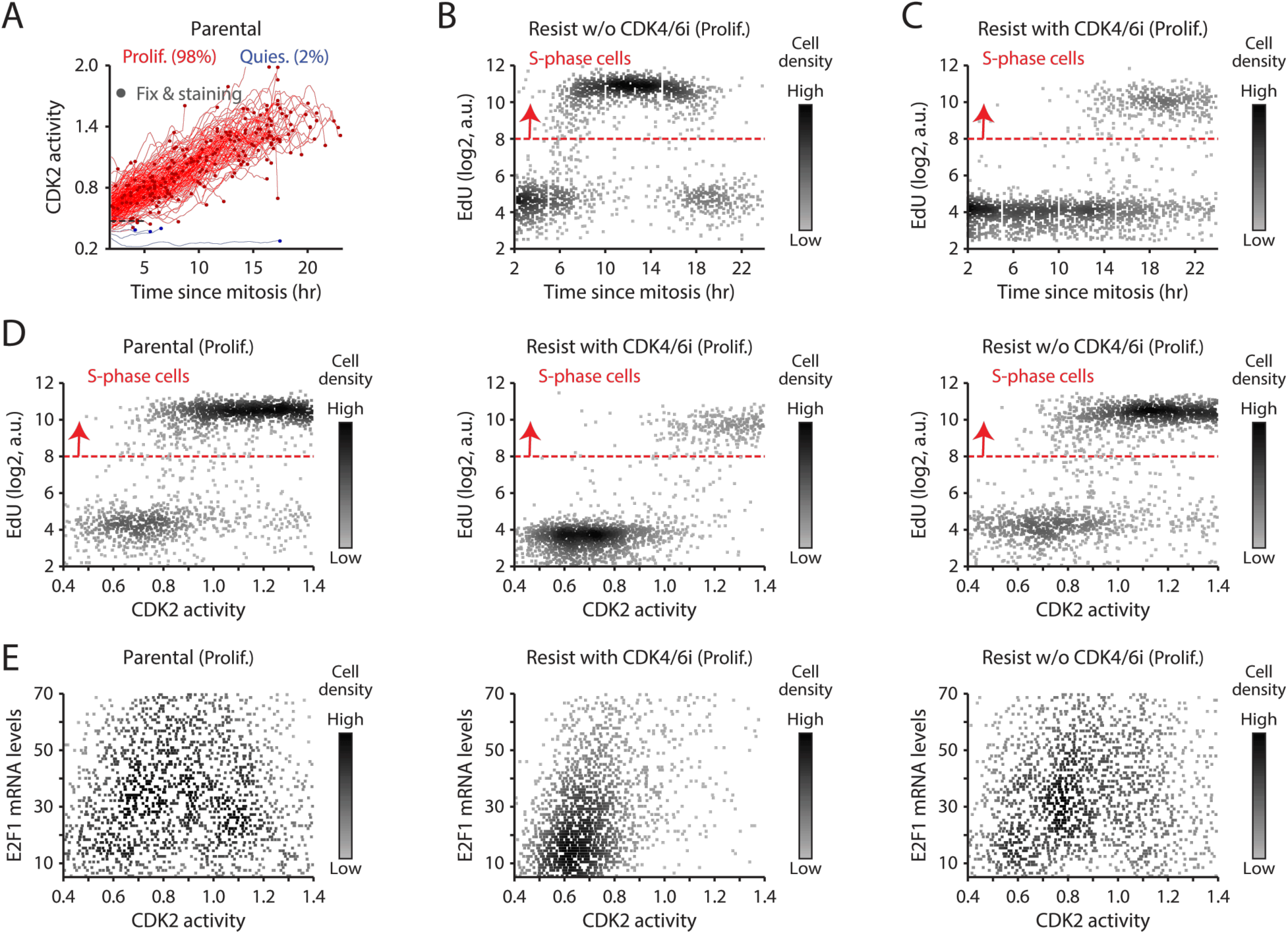
Slow and heterogeneous G1/S transition in drug-resistant cells maintained with CDK4/6i treatment. (A) Single-cell traces showing CDK2 activity aligned by mitosis in proliferating (red) and quiescent (blue) drug-naïve MCF-7 cells. Circles indicate the time of fixation and staining (*n* = 200 cells). (B, C) Scatterplot of EdU intensity against time since mitosis in drug-resistant cells without (B) and with (C) continuous palbociclib (1 µM) treatment. Red dotted line indicates the S-phase threshold (*n* = 2,000 cells/condition). (D, E) Scatterplot of EdU intensity (D) and E2F1 mRNA levels (E) against CDK2 activity (*n* = 2,000 cells/condition).

**Supplementary Figure 6.**
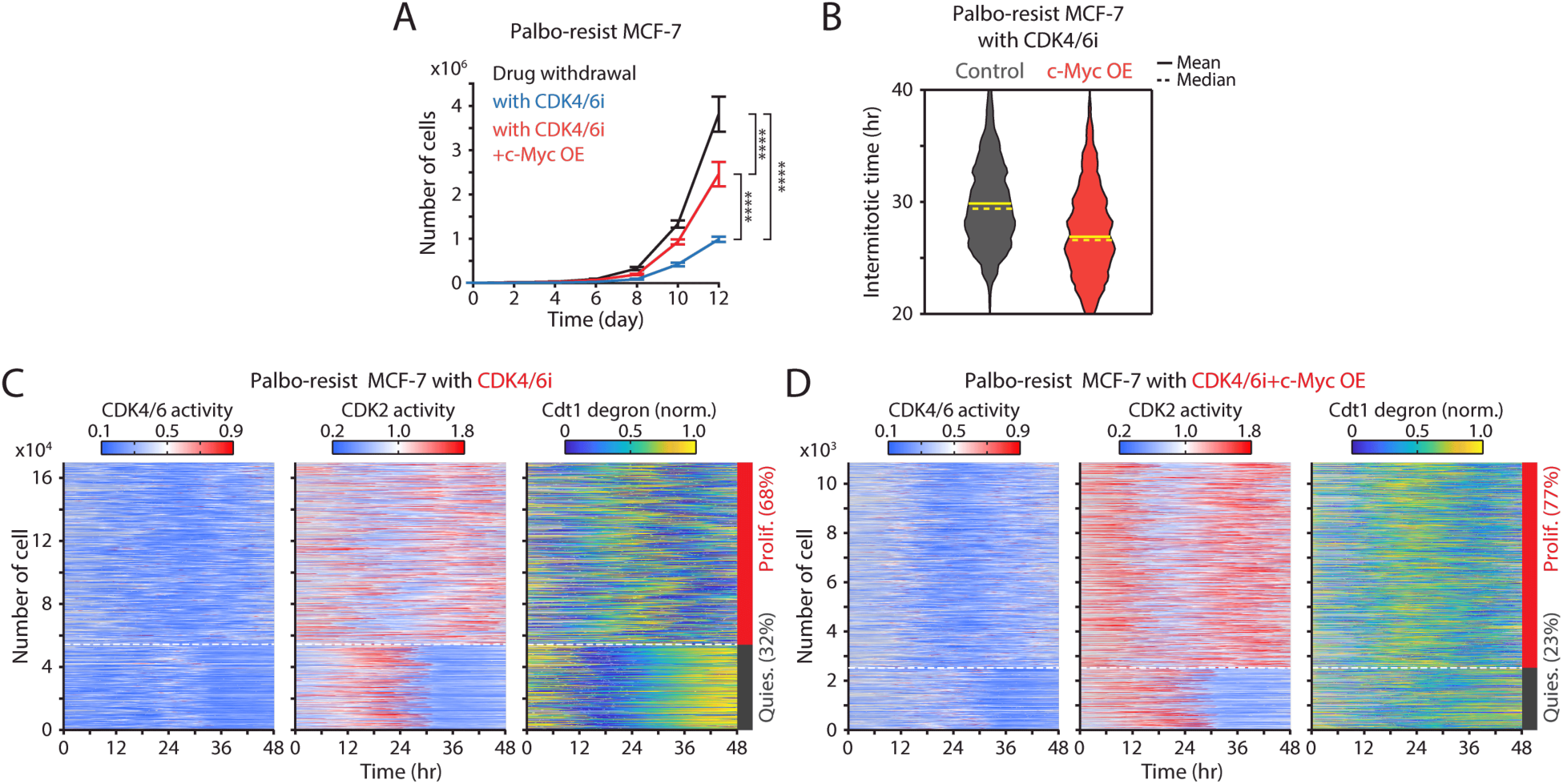
c-Myc overexpression facilitates CDK4/6i resistance by accelerating cell-cycle progression. (A) Growth curves of drug-resistant cells with treatment discontinuation or continuous palbociclib (1 µM) treatment, without or with c-Myc overexpression. Data represent mean ± SD (*n* = 3 biological replicates). Statistical significance was determined using two-way ANOVA with Tukey’s post-hoc analysis (**** *P* < 0.0001). (B) Intermitotic time of drug-resistant cells without and with c-Myc overexpression. Solid and dashed yellow lines represent mean and median, respectively (*n* > 4,500 cells/condition). (C, D) Heatmaps of single-cell traces for CDK4/6 (left) and CDK2 (middle) activities, and Cdt1-degron intensity (right) in drug-resistant cells undergoing continuous palbociclib (1 µM) treatment without (C) and with (D) c-Myc overexpression. Proliferating cells were identified based on CDK2 activity (>1 for more than 2 hr between 30 and 48 hr).

**Supplementary Figure 7.**
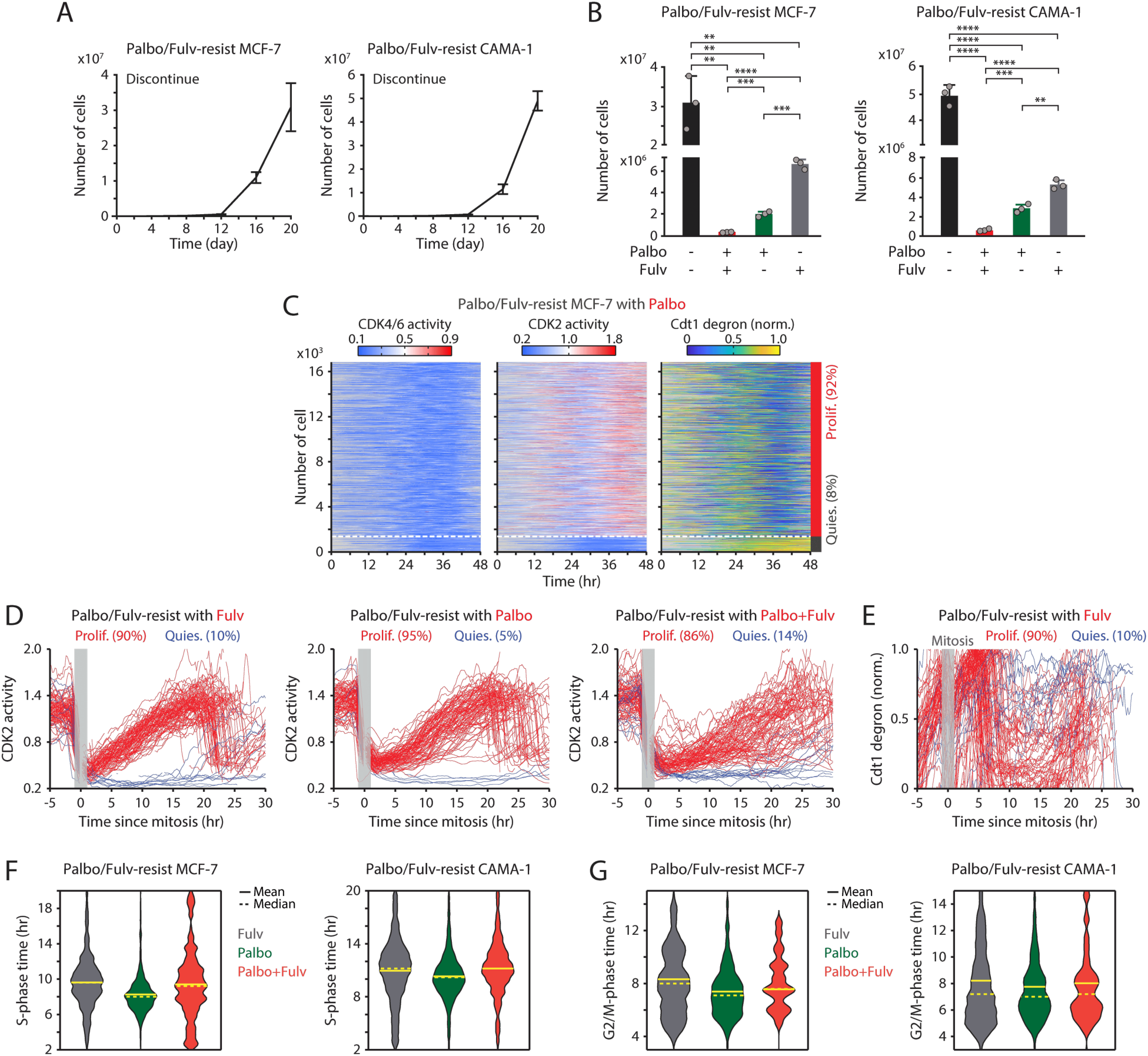
Maintaining CDK4/6i and ET synergistically suppresses the growth of drug-resistant cells. (A) Growth curves of MCF-7 (left) and CAMA-1 (right) cells resistant to palbociclib and fulvestrant withdrawn from drug treatment. Data represent the mean ± SD (*n* = 3 biological replicates). (B) Cell numbers 20 days after treatment with DMSO, palbociclib (1 µM), fulvestrant (500 nM), or their combination. Data represent mean ± SD (*n* = 3 biological replicates). Statistical significance was determined with an unpaired *t*-test (** *P* < 0.01, *** *P* < 0.001, **** *P* < 0.0001). (C) Heatmaps of single-cell traces for CDK4/6 (left) and CDK2 (middle) activities, and Cdt1-degron intensity (right) in combination drug-resistant cells maintained with palbociclib (1 µM). Proliferating cells were identified based on CDK2 activity (>1 for more than 2 hr between 30 and 48 hr). (D) Single-cell traces of CDK2 activity aligned by mitosis in combination drug-resistant cells treated with continuous fulvestrant (500 nM) (left), palbociclib (1 µM) (middle), or their combination (right). Based on CDK2 activity, cells were classified into proliferation (red) or quiescence (blue). The time of mitosis is marked in gray. (E) Single-cell traces of Cdt1-degron intensity aligned by mitosis in combination drug-resistant cells treated with continuous fulvestrant (500 nM) alone. (F, G) Violin plots showing S-phase duration (*n* > 200 cells/condition) (F) and G2/M-phase duration (*n* > 20 cells/condition) (G) in MCF-7 (left) and CAMA-1 (right) cells. Solid and dashed yellow lines indicate mean and median, respectively.

**Supplementary Figure 8.**
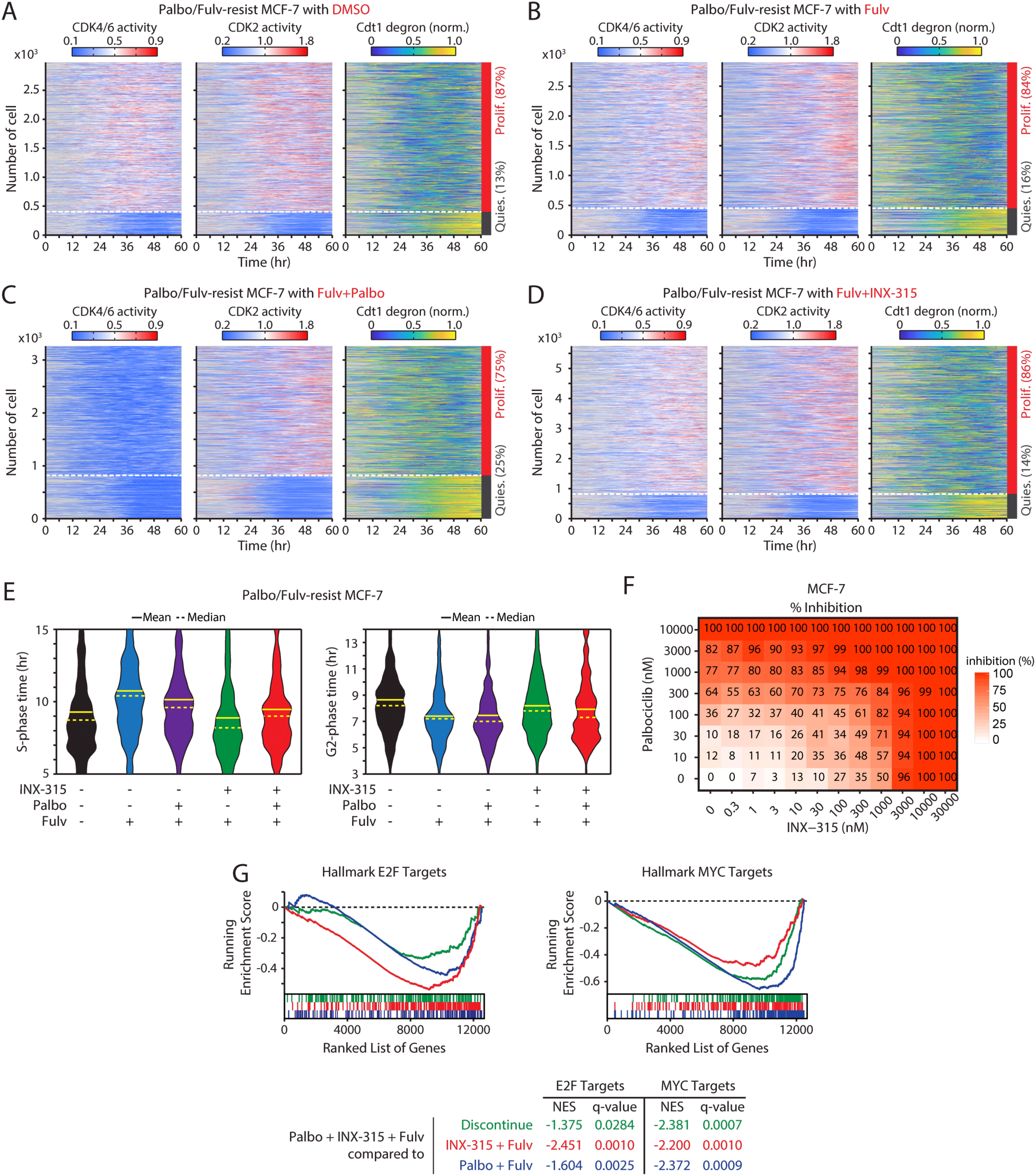
Combining CDK2i with CDK4/6i and ET effectively suppresses the growth of drug-resistant cells. (A–D) Heatmaps of single-cell traces for CDK4/6 (left) and CDK2 (middle) activities, and Cdt1-degron intensity (right) in drug-resistant cells under various conditions: treatment discontinuation (A), fulvestrant (500 nM) alone (B), fulvestrant + palbociclib (1 µM) (C), or fulvestrant + INX-315 (100 nM) (D) over one week prior to imaging. Proliferating cells were identified based on CDK2 activity (>1 for more than 2 hr between 30 and 48 hr). (E) Violin plots showing S-phase duration (*n* > 400 cells/condition) (left) and G2/M-phase duration (*n* > 150 cells/condition) (right) in drug-resistant MCF-7 cells treated with the indicated drug. Solid and dashed yellow lines indicate mean and median, respectively. (F) Two-dimensional titration of palbociclib (0–10 µM) and INX-315 (0–30 µM) in MCF-7 cells for 48 hr (*n* = 3 biological replicates). (G) GSEA plots for hallmark E2F (left) and MYC (right) target genes in palbociclib/fulvestrant-resistant MCF-7 cells treated with the triple combination, compared to other drug conditions.

**Supplementary Figure 9.**
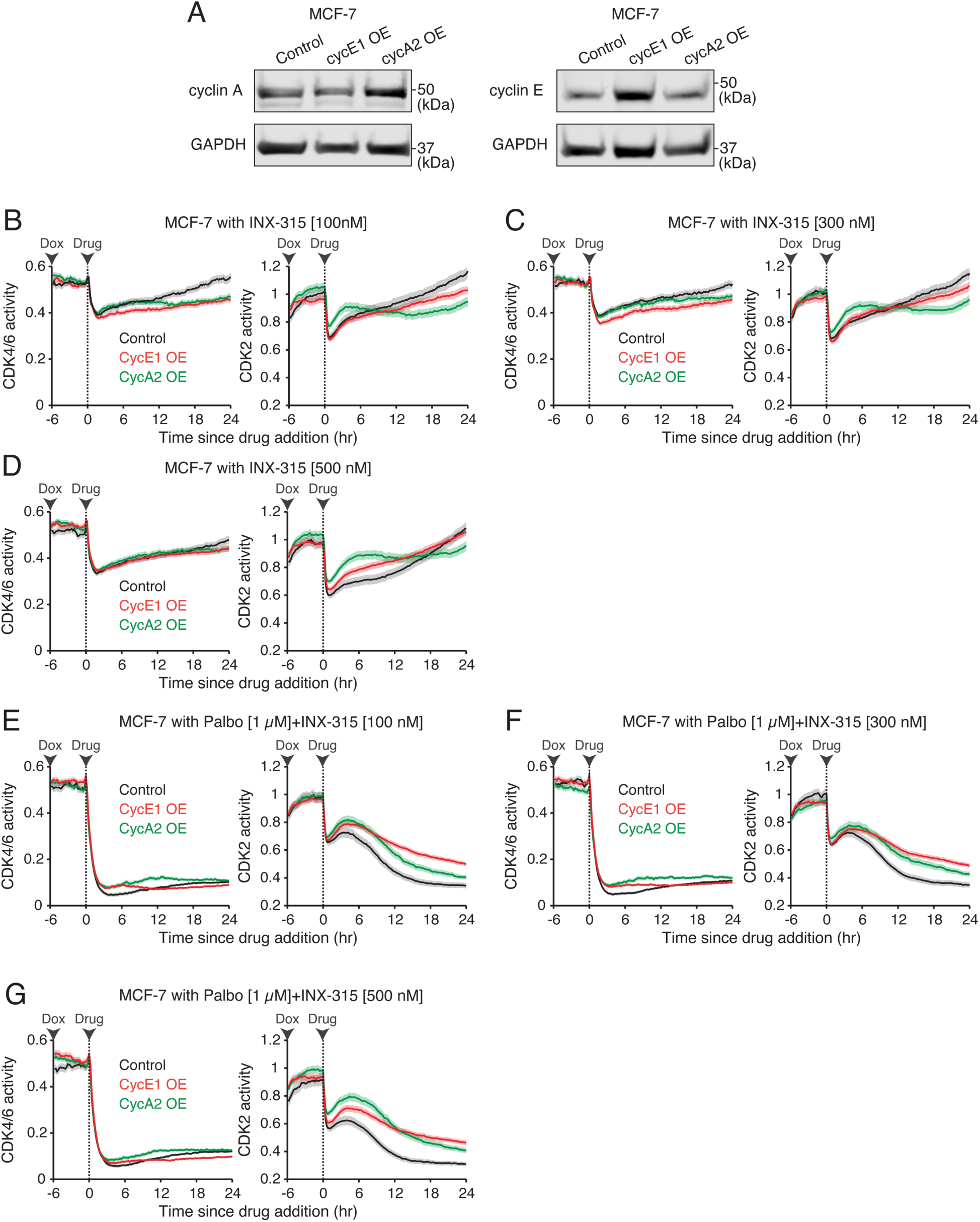
Cyclin E or A overexpression attenuates full CDK2 inhibition by CDK4/6i and CDK2i combination. (A) Immunoblot showing the expression of GAPDH with cyclin A (left) or cyclin E (right) in MCF-7 cells with or without cyclin E1 or A2 overexpression. Cells were treated with doxycycline (500 nM) for 24 hr. (B–G) Averaged traces of CDK4/6 (left) and CDK2 (right) activities in MCF-7 cells with or without cyclin E1 and A2 overexpression. Cells were treated with indicated doxycycline (500 nM) 6 hr before the addition of indicated drugs. Data represent mean ± 95% confidence intervals (*n* > 850 cells/condition).

**Supplementary Figure 10.**
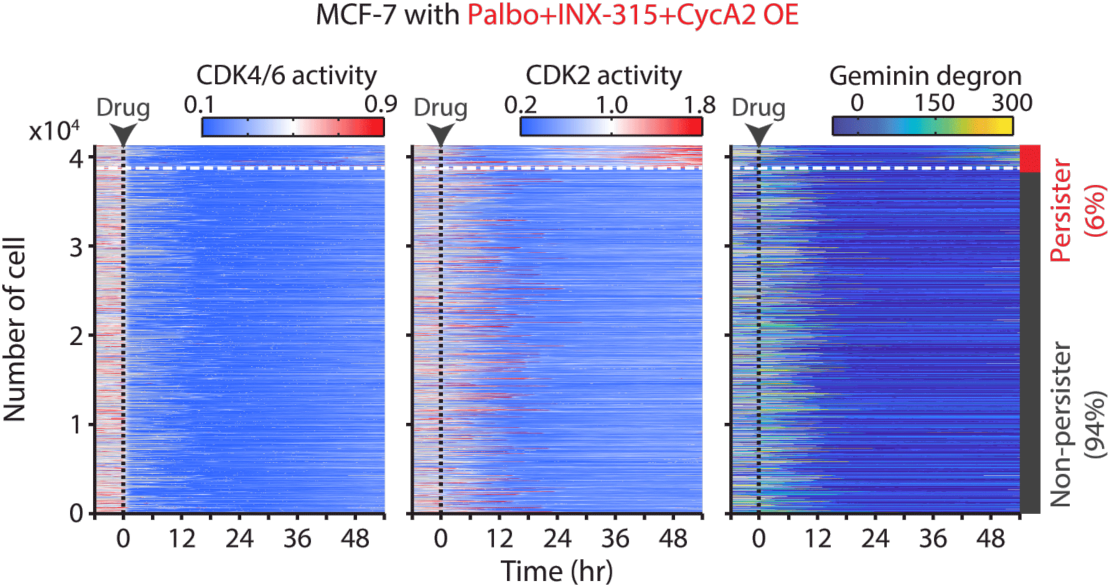
Cyclin E overexpression facilitates adaptation to CDK4/6i and CDK2i combination. Heatmaps of single-cell traces for CDK4/6 (left) and CDK2 (middle) activities, and Geminin-degron intensity (right) in drug-naïve MCF-7 cells overexpressing cyclin A and treated with palbociclib (1 µM) and INX-315 (100 nM). Proliferating cells were identified based on CDK2 activity (>1 for more than 2 hr between 30 and 48 hr).

## Materials and Methods

### Cell culture

MCF-7 (ATCC, HTB-22), CAMA-1 (ATCC, HTB-21), and MDA-MB-231 (ATCC, CRM-HTB-26) cells were cultured in Dulbecco’s Modified Eagle Medium (DMEM; Genesee Scientific, 25-500) supplemented with 10% fetal bovine serum (FBS; Gibco, A3160601). T47D (ATCC, HTB-133) cells were maintained in RPMI-1640 (Genesee Scientific, 25-206) with 10% FBS. All cell lines were grown at 37 °C in a humidified incubator with 5% CO₂ and were routinely tested and confirmed negative for mycoplasma contamination.

### Drugs and Chemicals

For in vitro experiments, palbociclib (Selleck Chemicals, S1116), INX-315 (MedChemExpress, HY-162001), and fulvestrant (Selleck Chemicals, S1191) were dissolved in DMSO (Sigma-Aldrich, D2438). For in vivo studies, palbociclib (MedChemExpress, HY-50767), ribociclib (MedChemExpress, HY-15777), and abemaciclib (MedChemExpress, HY-16297A) were prepared in corn oil (Spectrum Chemical, CO136) containing 10% DMSO. 5-Ethynyl-2′-deoxyuridine (EdU; Sigma-Aldrich, 900584) and AFDye-647 picolyl azide (Click Chemistry Tools, 1300) were used to label DNA synthesis.

### Antibodies

Primary antibodies were from Cell Signaling Technology: Rb (9309), phospho-Rb (Ser807/811; 8516), and c-Myc (5605); from Abcam: CDK6 (ab124821), cyclin E (ab32103), and cyclin A (ab32386); and from BD Biosciences: Rb (554136). The BD Biosciences Rb antibody was used for Rb visualization in cells expressing the CDK4/6 sensor. Secondary antibodies for immunofluorescence were Alexa Fluor 488 goat anti-mouse (Thermo Fisher Scientific, A32723) and Alexa Fluor 568 goat anti-rabbit (Thermo Fisher Scientific, A11036). For immunoblotting, IRDye secondaries were used: goat anti-mouse 800CW (LI-COR, 926-32210) and goat anti-rabbit 680RD (LI-COR, 926-68071).

### DNA constructs and cell line generation

DNA constructs were generated as described previously (8,35,42,43,74). Briefly, Gibson assembly was used to clone H2B-iRFP670-p2a-mCerulean-Cdt1 (1–100) (Addgene, 223965), H2B-iRFP670-p2a-mCerulean-Geminin (1–110) (Addgene, 223959), and DHB (995–1087)-mVenus-p2a-mCherry-Rb (886–928) (Addgene, 126679) into pLenti-IRES vectors bearing puromycin, blasticidin, or neomycin selection markers. Doxycycline-inducible c-Myc, cyclin E1, and cyclin A2 constructs were generated by inserting PCR products into pCW57.1 (Addgene, 50661) after NheI/BamHI digestion.

Stable cell lines were created by lentiviral transduction. Lentiviral plasmids and packaging vectors pMDLg/pRRE (Addgene, 12251), pRSV-Rev (Addgene, 12253), and pCMV-VSV-G (Addgene, 8454) were transfected into HEK-293T cells using polyethyleneimine in Opti-MEM (Thermo Fisher Scientific, 31985070). Viral supernatants were collected at 72 and 96 h, pooled, clarified (1,200 rpm, 5 min), filtered (0.45 µm; Millipore, SLHA033SB), concentrated (Amicon Ultra-15; Millipore, UFC910024; 4,000 rpm, 10 min), and stored at −80 °C. Target cells were infected in the presence of polybrene (5 µM) for 48 h and selected with puromycin (1 µg/mL; InvivoGen, ant-pr), blasticidin (10 µg/mL; InvivoGen, ant-bl), or neomycin (800 µg/mL; Thermo Fisher Scientific, BP673-5), or by FACS based on introduced fluorescent reporters.

### CDK6 and Rb knockout cell lines

Rb knockout in MCF-7 cells was previously described and validated (42). For CDK6 knockout, MCF-7 cells were transfected with crRNA:tracrRNA-ATTO550 duplexes complexed with recombinant Cas9 to form RNPs. ATTO550-positive cells were single-cell sorted by flow cytometry, expanded, and screened by immunoblotting. Two gRNAs were used to target CDK6:

- CDK6 gRNA 1: 5’-GACCACGUUGGGGUGCUCGAGUUUUAGAGCUAUGCU-3’
- CDK6 gRNA 2: 5’-CUGGACUGGAGCAAGACUUCGUUUUAGAGCUAUGCU-3’

### Drug-resistant cell lines

To generate CDK4/6i resistance, breast cancer cell lines were continuously exposed to palbociclib (1 µM) for over a month, with media and drug refreshed every 2–3 days. To establish double resistance to palbociclib and fulvestrant, palbociclib-resistant cells were then treated with fulvestrant (500 nM) for >2 months under the same refresh schedule. Resistance was confirmed by drug titration and determination of the half-maximal inhibitory concentration, calculated from the fraction of S-phase cells relative to drug-naïve controls.

### Live-cell reporhters

H2B-iRFP670 was used to segment nuclei and track single cells. A fluorescent Cdt1 degron (aa 1–100) (37) reported cell-cycle phase transitions; fluorescent geminin was used to mark S-phase entry. Kinase translocation reporters (KTRs) for CDK4/6 and CDK2 (35,36) monitored kinase-dependent phosphorylation of specific substrates (Rb C-terminus [aa 886–928] for CDK4/6 and DNA helicase B [DHB; aa 994–1087] for CDK2), driving regulated nucleo-cytoplasmic shuttling. The cytoplasmic/nuclear fluorescence intensity ratio provided a quantitative readout of kinase activity. Because the CDK4/6 KTR contains a degenerate CDK2 motif and partially reports CDK2 activity in S/G2, we applied a linear-regression–derived correction based on the CDK2 reporter (35,42,44,74,75):

- MCF-7: CDK4/6 activity = (CDK4/6 reporter) − 0.41 × (CDK2 reporter)
- MDA-MB-231: CDK4/6 activity = (CDK4/6 reporter) − 0.35 × (CDK2 reporter)

### Immunofluorescence and mRNA fluorescence in situ hybridization (FISH)

Cells were seeded in glass-bottom 96 well plates (Cellvis, P96-1.5H-N) at least 16 hr before experiments. S-phase fraction was assessed by EdU incorporation. Cells were pulsed with EdU (10 µM) for 15 min at 37°C, fixed in 4% formaldehyde (Thermo Scientific, 28906) in PBS supplemented with 10 mM HEPES (Sigma-Aldrich, H3537) for 15 min at room temperature (RT), and permeabilized in 0.2% Triton X-100 in PBS (PBS-T; Sigma-Aldrich, T8787) for 15 min. EdU was detected by click chemistry (15 min at RT) in 2 mM CuSO4 (Sigma-Aldrich, C1297), 20 mg/ml sodium ascorbate (Sigma-Aldrich, A4034), and 3 µM Alexa Fluor Dye 647 picolyl azide (Click Chemistry Tool, 1300) prepared in TBS (pH 8.3; Sigma-Aldrich, T6066).

For immunofluorescence (IF), cells were blocked (1 hr, RT) in PBS containing 10% FBS, 1% BSA, 0.1% Triton X-100, and 0.01% NaN3. Primary antibodies were incubated overnight at 4 °C in blocking buffer: anti-phospho-Rb (Ser807/811; Cell Signaling Technology, 8516), anti-Rb (BD Biosciences, 554136), and anti-c-Myc (Cell Signaling Technology, 5605). The next day, Alexa Fluor 488- or 568-conjugated secondary antibodies (Thermo Fisher Scientific) were applied for 1 hr at RT (1:2,000 in blocking buffer). Nuclei were counterstained with Hoechst 33342 (Thermo Fisher Scientific, 62249; 10 mg/mL stock diluted 1:10,000 in PBS, 15 min, RT). Cells were washed with PBS between steps and stored in PBS until imaging. For mRNA FISH, the ViewRNA ISH Cell Assay Kit (Thermo Fisher Scientific, QVC0001) and an E2F1 probe set (Thermo Fisher Scientific, VA1-12108-VC) were used according to the manufacturer’s instructions.

For experiments in Figure 3, we used a live–fixed pipeline. Asynchronously proliferating cells were imaged live for ≥48 h (37 °C, 5% CO₂; acquisition parameters detailed in “Live and fixed cell image acquisition”). Histone H2B fluorescence was used to segment/track nuclei and define time since mitosis (t = 0 at anaphase), and a CDK2 KTR provided CDK2 activity in the last live frame. Immediately after the live acquisition, the same wells were pulsed with EdU (10 µM, 15 min) and fixed/permeabilized as above. To prevent interference from fluorescent proteins during fixed assays, we photobleached residual fluorescence with 3% H₂O₂ + 20 mM HCl in PBS for 2 hr at RT. We then performed click-chemistry EdU detection, IF for phospho-Rb (Ser807/811) and total Rb, and RNA FISH for E2F1.

### Growth curve measurement

To assess population growth, 5,000 cells were seeded per well in 24-well plates (Thermo Fisher Scientific, FB012929). At the indicated time points, cells were harvested by trypsinization, resuspended in 1 mL growth medium, and counted manually with a hemocytometer (Hausser Scientific, HS-3510). Cells were replated as needed for subsequent time points.

### Live-cell, fixed-cell, and tumor tissue image acquisition

Cells were maintained at 30–80% confluency in 96-well glass-bottom plates (Cellvis, P96-1.5P) (75). Phenol red-free media were used to minimize background fluorescence. Imaging was performed on an inverted Nikon Eclipse Ti-2 microscope equipped with a Hamamatsu ORCA-Fusion camera. Objectives were 10× (Nikon CFI Plan Apo Lambda, NA 0.45; no binning) or 20× (Nikon CFI Plan Apo Lambda, NA 0.75; 2×2 binning). Excitation/emission settings were: CFP, 440/482 nm; GFP, 488/525 nm; YFP, 514/535 nm; mCherry, 594/609 nm; Cy5, 640/690 nm. Images were acquired using NIS-Elements AR v5.21.03. Live-cell imaging was performed in a humidified 37 °C/5% CO₂ chamber, acquiring 3 sites per well every 12 min. For fixed-cell imaging, 9 (10×) or 32 (20×) sites per well were collected. Total exposure time was kept <500 ms per time point. mRNA FISH images were collected with 1.5 µm z-stacks. Whole-tumor cross-sections were imaged on a Zeiss Axio Observer 7 with Apotome 2 and a Hamamatsu ORCA-Flash 4 camera using a 10× Zeiss Plan-Apochromat objective (NA 0.45; no binning) at 1.5 µm z-stack intervals.

### Image processing and analysis

Live- and fixed-cell images were processed using custom MATLAB scripts (MathWorks, R2021a). Flat-field correction was applied to reduce illumination bias. For fixed-cell analysis, nuclei were segmented on Hoechst using a histogram-curvature threshold. For live-cell analysis, nuclei were segmented using H2B-iRFP670 with a Laplacian-of-Gaussian blob detector; adjacent nuclei were separated by marker-based watershed. Background was corrected by subtracting the 50th percentile of non-nuclear pixels. For live-to-fixed alignment, cells were segmented in the final live frame and matched to the corresponding fixed-cell field. Cell tracks were generated with a deflection-bridging algorithm. Mitoses were identified by the appearance of two daughter nuclei adjacent to a prior nucleus and by H2B intensity (each daughter about 45–55% of the mother). Cytoplasmic signal was quantified as the median intensity within a ring 0.65–3.25 µm outside the nuclear boundary; cells with overlapping rings were excluded. For RNA FISH, whole-cell areas were estimated by dilating nuclear masks by 50 µm; cells overlapping neighbors were excluded. Raw FISH images were processed with a top-hat filter (4 µm circular kernel) to generate puncta masks. Tissue images were analyzed in ImageJ.

### Immunoblot

Cells were rinsed with ice-cold PBS and lysed in 300 µL CHAPS buffer (50 mM Tris-HCl, 150 mM NaCl, 1 mM EDTA, 10 mM N-ethylmaleimide, 0.3% CHAPS, 1 mM PMSF) supplemented with Halt Protease Inhibitor (Thermo Fisher Scientific, 1861279) and phosphatase inhibitor (Roche, 4906845001) for 5 min on ice. Lysates were cleared (14,000 × g, 10 min, 4 °C). Protein concentration was determined with the Pierce 660 nm assay (Thermo Fisher Scientific, 22660). Samples (12 µg) were denatured in LDS buffer (Thermo Fisher Scientific, NP0007) at 70 °C for 10 min, resolved on NuPAGE 4–12% Bis-Tris gels (Thermo Fisher Scientific, NP0322BOX), and transferred to 0.45 µm PVDF membranes (Bio-Rad, 1620174) using a Trans-Blot Turbo (Bio-Rad, 1704150) with 1× transfer buffer (Bio-Rad, 10026938). Membranes were blocked (LI-COR Intercept, 927-60001, 1 hr, RT) and incubated overnight (4 °C) with primary antibodies—Rb (CST, 9309), phospho-Rb Ser807/811 (CST, 8516), CDK6 (Abcam, ab124821), or β-actin (CST, 3700S)—diluted 1:1,000 in blocking buffer. After TBS-T washes (20 mM Tris, 150 mM NaCl, 0.1% Tween-20, pH 7.5), membranes were incubated with IRDye 800CW goat anti-mouse (LI-COR, 926-32210) or IRDye 680RD goat anti-rabbit (LI-COR, 926-68071) secondaries (1:2,000; 2 hr, RT). Blots were imaged on a LI-COR Odyssey and processed in Image Studio Lite v5.2.

### RNA sequencing

Total RNA was extracted using the GeneJET RNA Purification Kit (Thermo Fisher Scientific, K0732) according to the manufacturer’s protocol. After trypsinization and PBS wash, cells were lysed in 600 µL lysis buffer supplemented with β-mercaptoethanol (Bio-Rad, 1610710), homogenized, mixed with 360 µL 100% ethanol (Fisher Scientific, BP28184), and purified on spin columns. RNA was eluted in nuclease-free water (Thermo Fisher Scientific, 10977-015). Azenta Life Sciences performed library preparation and sequencing. Count-matrix analysis was conducted in R using DESeq2 (v1.44.0) (76) with log₂ fold-change shrinkage via apeglm (77). Gene sets (Hallmark, C2, C5) were obtained from MSigDB via msigdbr (v7.5.1). Heatmaps were generated with ComplexHeatmap (78). GSEA was performed with clusterProfiler (v4.12.6) using Hallmark annotations, a maximum gene-set size of 500, and 10,000 permutations; gene sets with nominal *P* < 0.05 and FDR < 0.25 were considered significant.

### Animal

All procedures were approved by the Institutional Animal Care and Use Committee at Columbia University Irving Medical Center and conducted in accordance with NIH guidelines. Female Foxn1-/- J:NU mice (6 weeks old; The Jackson Laboratory, 007850) were housed in a specific-pathogen-free barrier facility on a 12-hr light/dark cycle with ad libitum food and water and were monitored per institutional humane-endpoint policies.

### Xenograft experiments

Female J:NU mice were anesthetized by intraperitoneal injection of ketamine (45 mg/kg) plus xylazine (5 mg/kg). Drug-naïve MCF-7 cells (8 × 10⁶ per mouse) were resuspended 1:1 in PBS:Geltrex LDEV-Free Reduced Growth Factor Basement Membrane Matrix (Thermo Fisher Scientific, A1413201) and injected orthotopically into the abdominal mammary fat pad. Tumors were measured twice weekly with digital calipers; volumes were calculated as (width² × length) × 0.5. When tumors reached approximately 100 mm³, mice received daily palbociclib (50 mg/kg, oral gavage). Upon regrowth to 145–155 mm³, mice were randomized (n = 5/group) to: (1) treatment discontinuation; (2) palbociclib 50 mg/kg daily; (3) abemaciclib 50 mg/kg daily; or (4) ribociclib 200 mg/kg daily for 32 days. At study end, mice were anesthetized and perfused with 1% formaldehyde prior to tumor collection.

### Immunohistochemistry

After vascular perfusion with 1% formaldehyde in PBS, tumors were fixed in 1% formaldehyde (1 hr, 4 °C), washed in PBS-T, and cryoprotected in 30% sucrose (overnight, 4 °C). Samples were embedded in OCT (Thermo Fisher Scientific, 23-730-571), frozen at −80 °C, and sectioned at 40 µm onto Superfrost Plus slides (Thermo Fisher Scientific, 12-550-15). Sections were rehydrated in PBS-T, blocked in 5% normal goat serum (Jackson ImmunoResearch, 005-000-121) for 1 hr (RT), and incubated with recombinant anti-Rb (Abcam, ab181616; 1:1,000 in 5% serum/PBS-T) overnight (RT). After PBS-T washes, sections were incubated with Alexa Fluor 488 goat anti-rabbit (Thermo Fisher Scientific, A32731; 1:500 in PBS-T) for 4 hr (RT), counterstained with DAPI (Sigma-Aldrich, D9542; 1:500, 10 min), and mounted in Fluoromount-G (Invitrogen, 00-4958-02) for imaging.

### Statistics and Reproducibility

Statistical analyses were performed in GraphPad Prism v10.2.0. For parametric data, unpaired two-tailed Student’s t-tests or one-way ANOVA with Tukey’s post-hoc analysis were used as appropriate; paired two-tailed t-tests were used for paired comparisons. The specific tests and sample sizes are reported in the figure legends; a comprehensive summary is provided in Supplementary Table 1. All experiments were reproduced in at least two independent biological replicates.

## Data Availability

All data supporting the findings of this study are available within the article, the Source Data files, and Supplementary Table 1. Bulk RNA-seq data have been deposited in the Gene Expression Omnibus under accession GSE279160.

## Authors’ Disclosures

No disclosures were reported.

## Authors’ Contributions

(A) **J. Armand**: Conceptualization, investigation, data analysis, and resources; **S. Kim**: Investigation, data analysis, and resources; **K. Kim**: Investigation, data analysis, and resources; **E. Son**: Data analysis; **M. Kim**: Resources and funding acquisition; **K. Kalinsky**: Conceptualization; **H. Yang**: Conceptualization, investigation, data analysis, resources, writing the manuscript, and funding acquisition. All authors discussed and edited the manuscript.

## Acknowledgments

We thank Sarat Chandarlapaty for providing the CAMA-1 cells and Kevin Gardner for offering MDA-MB-231 cells. We thank Caitlin O’Neil for assisting with tissue imaging. We thank Anusha Shanabag for her assistance with the RNA-seq data processing. This work was supported by Neuroendocrine Tumor Research Foundation (M.K., 855538), Melanoma Research Foundation (M.K. and H.Y.), Research Scholar Grant (M.K., RSG-22-167-01-MM and H.Y., RSG-22-101-01-CDP), V Scholar (H.Y., V2023-017), NIH grants R37-CA266270 (M.K.), R03-AG073833 (M.K.), R01-GM145884 (H.Y.), and HICCC Grant P30-CA013696 (M.K. and H.Y.). These studies used the resources of HICCC Flow Cytometry Shared Resources (P30-CA013696).

## Notes

### Competing Interest Statement

The authors have declared no competing interest.

### Summary of Updates

The manuscript has been revised throughout according to the reviewers' comments. Added new figures: 3L, 3M, 6G, and S8F.

## References

1. Lei S, Zheng R, Zhang S, Wang S, Chen R, Sun K, et al. Global patterns of breast cancer incidence and mortality: A population-based cancer registry data analysis from 2000 to 2020. Cancer Commun (Lond) 2021;41:1183–94

2. Houghton SC, Hankinson SE. Cancer Progress and Priorities: Breast Cancer. Cancer Epidemiol Biomarkers Prev 2021;30:822–44

3. Sherr CJ, Beach D, Shapiro GI. Targeting CDK4 and CDK6: From Discovery to Therapy. Cancer Discov 2016;6:353–67

4. Watt AC, Goel S. Cellular mechanisms underlying response and resistance to CDK4/6 inhibitors in the treatment of hormone receptor-positive breast cancer. Breast Cancer Res 2022;24:17

5. Fassl A, Geng Y, Sicinski P. CDK4 and CDK6 kinases: From basic science to cancer therapy. Science 2022;375:eabc1495

6. Shanabag A, Armand J, Son E, Yang HW. Targeting CDK4/6 in breast cancer. Exp Mol Med 2025;57:312–22

7. Engeland K. Cell cycle regulation: p53-p21-RB signaling. Cell Death & Differentiation 2022;29:946–60

8. Kim S, Leong A, Kim M, Yang HW. CDK4/6 initiates Rb inactivation and CDK2 activity coordinates cell-cycle commitment and G1/S transition. Sci Rep 2022;12:16810

9. Fisher RP. Getting to S: CDK functions and targets on the path to cell-cycle commitment. F1000Research 2016;5:2374

10. Howlader N, Altekruse SF, Li CI, Chen VW, Clarke CA, Ries LA, et al. US incidence of breast cancer subtypes defined by joint hormone receptor and HER2 status. J Natl Cancer Inst 2014;106

11. Goel S, Bergholz JS, Zhao JJ. Targeting CDK4 and CDK6 in cancer. Nat Rev Cancer 2022;22:356–72

12. Hortobagyi GN, Stemmer SM, Burris HA, Yap YS, Sonke GS, Paluch-Shimon S, et al. Ribociclib as First-Line Therapy for HR-Positive, Advanced Breast Cancer. N Engl J Med 2016;375:1738–48

13. Johnston S, Martin M, Di Leo A, Im SA, Awada A, Forrester T, et al. MONARCH 3 final PFS: a randomized study of abemaciclib as initial therapy for advanced breast cancer. NPJ Breast Cancer 2019;5:5

14. Robertson JF, Howell A, Gorbunova VA, Watanabe T, Pienkowski T, Lichinitser MR. Sensitivity to further endocrine therapy is retained following progression on first-line fulvestrant. Breast Cancer Res Treat 2005;92:169–74

15. Vergote I, Robertson JF, Kleeberg U, Burton G, Osborne CK, Mauriac L. Postmenopausal women who progress on fulvestrant (’Faslodex’) remain sensitive to further endocrine therapy. Breast Cancer Res Treat 2003;79:207–11

16. Xie Y, Zhao Y, Gong C, Chen Z, Zhang Y, Zhao Y, et al. Treatment after Progression on Fulvestrant among Metastatic Breast Cancer Patients in Clinical Practice: a Multicenter, Retrospective Study. Scientific Reports 2019;9:1710

17. Steger GG, Bartsch R, Wenzel C, Pluschnig U, Hussian D, Sevelda U, et al. Fulvestrant (’Faslodex’) in pre-treated patients with advanced breast cancer: a single-centre experience. Eur J Cancer 2005;41:2655–61

18. Barrios C, Forbes JF, Jonat W, Conte P, Gradishar W, Buzdar A, et al. The sequential use of endocrine treatment for advanced breast cancer: where are we? Ann Oncol 2012;23:1378–86

19. Llombart-Cussac A, Harper-Wynne C, Perello A, Hennequin A, Fernandez-Ortega A, Colleoni M, et al. Second-Line Endocrine Therapy With or Without Palbociclib Rechallenge in Patients With Hormone Receptor-Positive/Human Epidermal Growth Factor Receptor 2-Negative Advanced Breast Cancer: PALMIRA Trial. J Clin Oncol 2025;43:2084–93

20. Mayer EL, Ren Y, Wagle N, Mahtani R, Ma C, DeMichele A, et al. PACE: A Randomized Phase II Study of Fulvestrant, Palbociclib, and Avelumab After Progression on Cyclin-Dependent Kinase 4/6 Inhibitor and Aromatase Inhibitor for Hormone Receptor-Positive/Human Epidermal Growth Factor Receptor-Negative Metastatic Breast Cancer. J Clin Oncol 2024;42:2050–60

21. Kalinsky K, Bianchini G, Hamilton E, Graff SL, Park KH, Jeselsohn R, et al. Abemaciclib Plus Fulvestrant in Advanced Breast Cancer After Progression on CDK4/6 Inhibition: Results From the Phase III postMONARCH Trial. J Clin Oncol 2025;43:1101–12

22. Jhaveri KL, Neven P, Casalnuovo ML, Kim SB, Tokunaga E, Aftimos P, et al. Imlunestrant with or without Abemaciclib in Advanced Breast Cancer. N Engl J Med 2024

23. Kalinsky K, Accordino MK, Chiuzan C, Mundi PS, Sakach E, Sathe C, et al. Randomized Phase II Trial of Endocrine Therapy With or Without Ribociclib After Progression on Cyclin-Dependent Kinase 4/6 Inhibition in Hormone Receptor–Positive, Human Epidermal Growth Factor Receptor 2–Negative Metastatic Breast Cancer: MAINTAIN Trial. Journal of Clinical Oncology 2023;41:4004–13

24. Martin JM, Handorf EA, Montero AJ, Goldstein LJ. Systemic Therapies Following Progression on First-line CDK4/6-inhibitor Treatment: Analysis of Real-world Data. Oncologist 2022;27:441–6

25. Ravani LV, Calomeni P, Vilbert M, Madeira T, Wang M, Deng D, et al. Efficacy of Subsequent Treatments After Disease Progression on CDK4/6 Inhibitors in Patients With Hormone Receptor-Positive Advanced Breast Cancer. JCO Oncol Pract 2025;21:832–42

26. Matson JP, Cook JG. Cell cycle proliferation decisions: the impact of single cell analyses. The FEBS Journal 2017;284:362–75

27. Pandey K, Park N, Park KS, Hur J, Cho YB, Kang M, et al. Combined CDK2 and CDK4/6 Inhibition Overcomes Palbociclib Resistance in Breast Cancer by Enhancing Senescence. Cancers (Basel) 2020;12

28. Freeman-Cook K, Hoffman RL, Miller N, Almaden J, Chionis J, Zhang Q, et al. Expanding control of the tumor cell cycle with a CDK2/4/6 inhibitor. Cancer Cell 2021;39:1404–21 e11

29. Dietrich C, Trub A, Ahn A, Taylor M, Ambani K, Chan KT, et al. INX-315, a selective CDK2 inhibitor, induces cell cycle arrest and senescence in solid tumors. Cancer Discov 2023

30. Al-Qasem AJ, Alves CL, Ehmsen S, Tuttolomondo M, Terp MG, Johansen LE, et al. Co-targeting CDK2 and CDK4/6 overcomes resistance to aromatase and CDK4/6 inhibitors in ER+ breast cancer. NPJ Precis Oncol 2022;6:68

31. Kudo R, Safonov A, Jones C, Moiso E, Dry JR, Shao H, et al. Long-term breast cancer response to CDK4/6 inhibition defined by TP53-mediated geroconversion. Cancer Cell 2024

32. Arora M, Moser J, Hoffman TE, Watts LP, Min M, Musteanu M, et al. Rapid adaptation to CDK2 inhibition exposes intrinsic cell-cycle plasticity. Cell 2023;186:2628–43 e21

33. Kumarasamy V, Wang J, Roti M, Wan Y, Dommer AP, Rosenheck H, et al. Discrete vulnerability to pharmacological CDK2 inhibition is governed by heterogeneity of the cancer cell cycle. Nature Communications 2025;16:1476

34. Dommer AP, Kumarasamy V, Wang J, O’Connor TN, Roti M, Mahan S, et al. Tumor Suppressors Condition Differential Responses to the Selective CDK2 Inhibitor BLU-222. Cancer Res 2025

35. Yang HW, Cappell SD, Jaimovich A, Liu C, Chung M, Daigh LH, et al. Stress-mediated exit to quiescence restricted by increasing persistence in CDK4/6 activation. Elife 2020;9

36. Spencer SL, Cappell SD, Tsai FC, Overton KW, Wang CL, Meyer T. The proliferation-quiescence decision is controlled by a bifurcation in CDK2 activity at mitotic exit. Cell 2013;155:369–83

37. Sakaue-Sawano A, Yo M, Komatsu N, Hiratsuka T, Kogure T, Hoshida T, et al. Genetically Encoded Tools for Optical Dissection of the Mammalian Cell Cycle. Mol Cell 2017;68:626–40 e5

38. Yang C, Li Z, Bhatt T, Dickler M, Giri D, Scaltriti M, et al. Acquired CDK6 amplification promotes breast cancer resistance to CDK4/6 inhibitors and loss of ER signaling and dependence. Oncogene 2017;36:2255–64

39. Li Q, Jiang B, Guo J, Shao H, Del Priore IS, Chang Q, et al. INK4 Tumor Suppressor Proteins Mediate Resistance to CDK4/6 Kinase Inhibitors. Cancer Discov 2022;12:356–71

40. Ji W, Zhang W, Wang X, Shi Y, Yang F, Xie H, et al. c-myc regulates the sensitivity of breast cancer cells to palbociclib via c-myc/miR-29b-3p/CDK6 axis. Cell Death Dis 2020;11:760

41. Wu X, Yang X, Xiong Y, Li R, Ito T, Ahmed TA, et al. Distinct CDK6 complexes determine tumor cell response to CDK4/6 inhibitors and degraders. Nat Cancer 2021;2:429–43

42. Kim S, Armand J, Safonov A, Zhang M, Soni RK, Schwartz G, et al. Sequential activation of E2F via Rb degradation and c-Myc drives resistance to CDK4/6 inhibitors in breast cancer. Cell Rep 2023;42:113198

43. Zhang M, Kim S, Yang HW. Non-canonical pathway for Rb inactivation and external signaling coordinate cell-cycle entry without CDK4/6 activity. Nature Communications 2023;14:7847

44. Kim S, Son E, Park HR, Kim M, Yang HW. Dual targeting CDK4/6 and CDK7 augments tumor response and anti-tumor immunity in breast cancer models. J Clin Invest 2025

45. Chung M, Liu C, Yang HW, Koberlin MS, Cappell SD, Meyer T. Transient Hysteresis in CDK4/6 Activity Underlies Passage of the Restriction Point in G1. Mol Cell 2019;76:562–73 e4

46. Johnson DG, Ohtani K, Nevins JR. Autoregulatory control of E2F1 expression in response to positive and negative regulators of cell cycle progression. Genes & Development 1994;8:1514–25

47. Yang HW, Chung M, Kudo T, Meyer T, Yang HW, Chung, Mingyu, Kudo T, et al. Competing memories of mitogen and p53 signalling control cell-cycle entry. Nature 2017;549:404–8

48. Westendorp B, Mokry M, Groot Koerkamp MJ, Holstege FC, Cuppen E, de Bruin A. E2F7 represses a network of oscillating cell cycle genes to control S-phase progression. Nucleic Acids Res 2012;40:3511–23

49. Shang Y, Hu X, DiRenzo J, Lazar MA, Brown M. Cofactor dynamics and sufficiency in estrogen receptor-regulated transcription. Cell 2000;103:843–52

50. Dhanyamraju PK, Schell TD, Amin S, Robertson GP. Drug-Tolerant Persister Cells in Cancer Therapy Resistance. Cancer Res 2022;82:2503–14

51. Sakaue-Sawano A, Kurokawa H, Morimura T, Hanyu A, Hama H, Osawa H, et al. Visualizing spatiotemporal dynamics of multicellular cell-cycle progression. Cell 2008;132:487–98

52. Peters J-M. The anaphase promoting complex/cyclosome: a machine designed to destroy. Nature Reviews Molecular Cell Biology 2006;7:644–56

53. Finn RS, Crown JP, Lang I, Boer K, Bondarenko IM, Kulyk SO, et al. The cyclin-dependent kinase 4/6 inhibitor palbociclib in combination with letrozole versus letrozole alone as first-line treatment of oestrogen receptor-positive, HER2-negative, advanced breast cancer (PALOMA-1/TRIO-18): a randomised phase 2 study. Lancet Oncol 2015;16:25–35

54. Finn RS, Martin M, Rugo HS, Jones S, Im S-A, Gelmon K, et al. Palbociclib and Letrozole in Advanced Breast Cancer. New England Journal of Medicine 2016;375:1925–36

55. Turner NC, Slamon DJ, Ro J, Bondarenko I, Im S-A, Masuda N, et al. Overall Survival with Palbociclib and Fulvestrant in Advanced Breast Cancer. New England Journal of Medicine 2018;379:1926–36

56. Dickler MN, Tolaney SM, Rugo HS, Cortés J, Diéras V, Patt D, et al. MONARCH 1, A Phase II Study of Abemaciclib, a CDK4 and CDK6 Inhibitor, as a Single Agent, in Patients with Refractory HR(+)/HER2(-) Metastatic Breast Cancer. Clin Cancer Res 2017;23:5218–24

57. Johnston S, Martin M, Di Leo A, Im S-A, Awada A, Forrester T, et al. MONARCH 3 final PFS: a randomized study of abemaciclib as initial therapy for advanced breast cancer. npj Breast Cancer 2019;5:5

58. Hortobagyi GN, Stemmer SM, Burris HA, Yap Y-S, Sonke GS, Hart L, et al. Overall Survival with Ribociclib plus Letrozole in Advanced Breast Cancer. New England Journal of Medicine 2022;386:942–50

59. Slamon DJ, Neven P, Chia S, Fasching PA, De Laurentiis M, Im S-A, et al. Overall Survival with Ribociclib plus Fulvestrant in Advanced Breast Cancer. New England Journal of Medicine 2019;382:514–24

60. Im S-A, Lu Y-S, Bardia A, Harbeck N, Colleoni M, Franke F, et al. Overall Survival with Ribociclib plus Endocrine Therapy in Breast Cancer. New England Journal of Medicine 2019;381:307–16

61. Tanaka S, Tak Y-S, Araki H. The role of CDK in the initiation step of DNA replication in eukaryotes. Cell Division 2007;2:16

62. Krude T, Jackman M, Pines J, Laskey RA. Cyclin/Cdk-dependent initiation of DNA replication in a human cell-free system. Cell 1997;88:109–19

63. Hafner M, Mills CE, Subramanian K, Chen C, Chung M, Boswell SA, et al. Multiomics Profiling Establishes the Polypharmacology of FDA-Approved CDK4/6 Inhibitors and the Potential for Differential Clinical Activity. Cell Chem Biol 2019;26:1067–80.e8

64. Navarro-Yepes J, Kettner NM, Rao X, Bishop CS, Bui TN, Wingate HF, et al. Abemaciclib Is Effective in Palbociclib-Resistant Hormone Receptor-Positive Metastatic Breast Cancers. Cancer Res 2023;83:3264–83

65. O’Leary B, Cutts RJ, Liu Y, Hrebien S, Huang X, Fenwick K, et al. The Genetic Landscape and Clonal Evolution of Breast Cancer Resistance to Palbociclib plus Fulvestrant in the PALOMA-3 Trial. Cancer Discov 2018;8:1390–403

66. Mao P, Cohen O, Kowalski KJ, Kusiel JG, Buendia-Buendia JE, Cuoco MS, et al. Acquired FGFR and FGF Alterations Confer Resistance to Estrogen Receptor (ER) Targeted Therapy in ER(+) Metastatic Breast Cancer. Clin Cancer Res 2020;26:5974–89

67. Wander SA, Cohen O, Gong X, Johnson GN, Buendia-Buendia JE, Lloyd MR, et al. The Genomic Landscape of Intrinsic and Acquired Resistance to Cyclin-Dependent Kinase 4/6 Inhibitors in Patients with Hormone Receptor-Positive Metastatic Breast Cancer. Cancer Discov 2020;10:1174–93

68. Formisano L, Stauffer KM, Young CD, Bhola NE, Guerrero-Zotano AL, Jansen VM, et al. Association of FGFR1 with ERalpha Maintains Ligand-Independent ER Transcription and Mediates Resistance to Estrogen Deprivation in ER(+) Breast Cancer. Clin Cancer Res 2017;23:6138–50

69. Costa C, Wang Y, Ly A, Hosono Y, Murchie E, Walmsley CS, et al. PTEN Loss Mediates Clinical Cross-Resistance to CDK4/6 and PI3Kalpha Inhibitors in Breast Cancer. Cancer Discov 2020;10:72–85

70. Zhu J, Blenis J, Yuan J. Activation of PI3K/Akt and MAPK pathways regulates Myc-mediated transcription by phosphorylating and promoting the degradation of Mad1. Proc Natl Acad Sci U S A 2008;105:6584–9

71. Tsai WB, Aiba I, Long Y, Lin HK, Feun L, Savaraj N, et al. Activation of Ras/PI3K/ERK pathway induces c-Myc stabilization to upregulate argininosuccinate synthetase, leading to arginine deiminase resistance in melanoma cells. Cancer Res 2012;72:2622–33

72. Li Z, Razavi P, Li Q, Toy W, Liu B, Ping C, et al. Loss of the FAT1 Tumor Suppressor Promotes Resistance to CDK4/6 Inhibitors via the Hippo Pathway. Cancer Cell 2018;34:893–905 e8

73. Kim K, Armand J, Kim S, Yang HW. E2F activity determines mitosis versus whole-genome duplication in G2-arrested cells. Nat Commun 2025;16:6677

74. Kim S, Carvajal R, Kim M, Yang HW. Kinetics of RTK activation determine ERK reactivation and resistance to dual BRAF/MEK inhibition in melanoma. Cell Rep 2023;42:112570

75. Yang HW. Investigating Heterogeneous Cell-Cycle Progression Using Single-Cell Imaging Approaches. Methods Mol Biol 2024;2740:263–73

76. Love MI, Huber W, Anders S. Moderated estimation of fold change and dispersion for RNA-seq data with DESeq2. Genome Biology 2014;15:550

77. Zhu A, Ibrahim JG, Love MI. Heavy-tailed prior distributions for sequence count data: removing the noise and preserving large differences. Bioinformatics 2019;35:2084–92

78. Gu Z, Hubschmann D. Make Interactive Complex Heatmaps in R. Bioinformatics 2022;38:1460–2

